# Holistic genetic barcoding reveals a lineage tree of tissue macrophage development

**DOI:** 10.1101/2024.11.29.625985

**Authors:** Larissa Frank, Daniel Postrach, Maurice Langhinrichs, Thomas Höfer, Hans-Reimer Rodewald

**Author notes:** These authors contributed equally to this work (Larissa Frank, Daniel Postrach, Maurice Langhinrichs).

## Abstract

Tissue macrophages are crucial for organ development and homeostasis, yet the developmental routes leading to tissue macrophages remain controversial. By combining unbiased *Polylox* barcoding with computational inference, we comprehensively mapped tissue macrophage ontogeny in mice from gastrulation to adult organs. Our data reveal a lineage tree of all tissue macrophages. This tree originates from a pan-hematopoietic progenitor at embryonic day (E)6.5 and branches, via oligolineage progenitors (E7.5-E9.5), into organ-specific macrophage lineages by E10.5. Barcode analysis in embryonic tissues and adult organ fragments suggests that local macrophage colonies, formed by organ-specific progenitors, persist into adulthood. Indeed, spatial fate mapping with *Polytope* barcoding detects local Kupffer cell colonies of the size predicted by the lineage tree model. Our findings uncover the origin of tissue-resident macrophages from a lineage tree, and show how holistic barcoding can comprehensively map complex cellular lineage specification.

**One-Sentence Summary:** Holistic, time- and space-resolved barcoding in the embryo reveals a unified lineage tree for all tissue macrophages, generating local macrophage colonies persisting in adult organs.

---

Tissue-specific macrophages are key components of tissue development and homeostasis (*1–3*). Their ontogeny has been intensively investigated by fate mapping in mice, yet no consensus has emerged. On the one hand, an early-arising (at E8.5) erythro-myeloid progenitor (EMP) has been suggested as a common source of all tissue macrophage populations in the adult mouse (*4–6*). On the other hand, distinct developmental waves of macrophage development have been invoked. In particular, it has been proposed that an early EMP generates the first wave that gives rise to microglia in the adult, whereas other adult tissue macrophages develop later, at about E13.5, via monocytes in the fetal liver (*7–10*). Another model posits that most tissue-resident macrophages, except microglia, are derived from fetal hematopoietic stem cells (*11*). Collectively, these data raise the fundamental question whether different kinds of tissue macrophages are generated by independent progenitors arising in distinct developmental stages or whether there is a single lineage tree of macrophage diversification deriving from a common macrophage progenitor.

Fate-mapping studies on macrophage ontogeny have used a wide range of gene loci to mark different progenitors via constitutive or inducible expression of Cre recombinase (including *Csf1r*, *Cx3cr1*, *Kit*, *Runx1*, *Tie2*, *Tnfrsf11a*, *S100a4*) (*4*, *5*, *9*, *11–14*). While providing physiological precursor-product relations, specific Cre drivers may be limited in their information, as it is difficult to ascertain whether and to what extent their expression covers in full the relevant progenitors at the appropriate developmental stages. Hence, contradictory results on the development of tissue macrophages could, at least in part, have resulted from different Cre drivers used. To address this major problem, we developed a holistic and quantitative approach for mapping cell ontogeny over developmental time.

## Resolving macrophage origin with unbiased progenitor barcoding

To map macrophage ontogeny, we non-invasively barcoded cells in the entire embryo at successive developmental stages and analyzed barcodes in mature tissue-resident macrophages and hematopoietic lineages. First, we induced *Polylox* DNA barcodes (*15*, *16*) at E6.5 in the embryo by applying 4-hydroxytamoxifen (OHT) (Fig. 1A, Fig. S1A) to pregnant mice carrying *Rosa^Polylox/CreERT2^* embryos, which acts for ∼1 day to generate barcodes (*17*). Barcodes were analyzed in adult mice in macrophage populations from the brain (microglia, M), liver (Kupffer cells, K), spleen (red pulp macrophages, R), lung (alveolar macrophages, A), kidney, and the peritoneum. Additionally, barcodes were examined in adult hematopoietic lineages (hematopoietic stem and progenitor cells (H), granulocytes, monocytes, lymphocytes) (Fig. 1A; Fig. S2, S3). Barcoding was comprehensive and uniform across cell populations, with >60% and >95% of *Polylox* reads in the two analyzed mice corresponding to recombined barcodes (i.e., *Polylox* loci with at least one Cre-mediated recombination event) (Fig. S1B). Hence, most progenitors that gave rise to the analyzed mature populations had been barcoded in the embryo.

**Figure 1.**
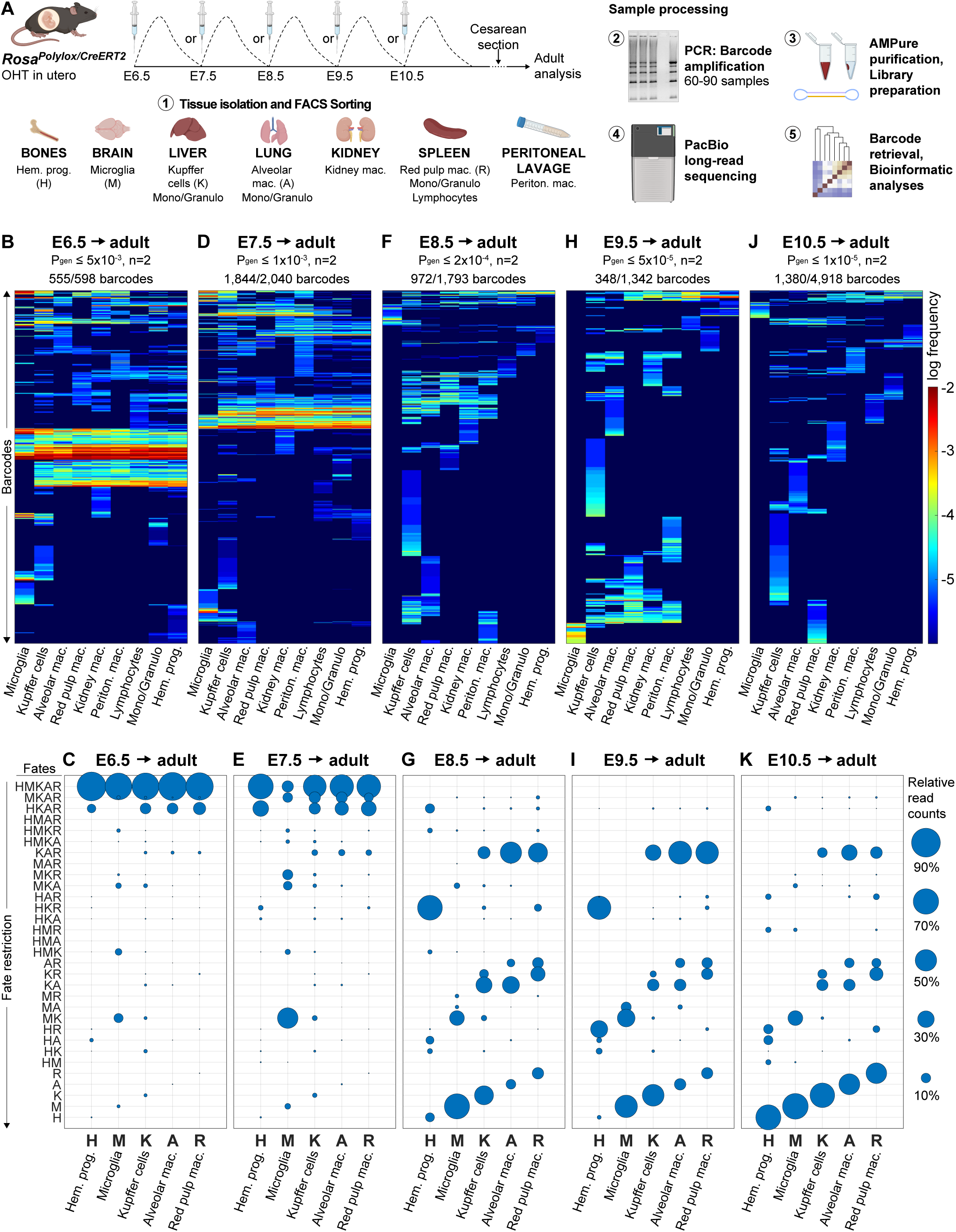
Unbiased *Polylox* barcoding *in utero* to trace progenitor fates into adult tissue macrophages and hematopoietic lineages. (**A**) Barcoding of *Rosa^Polylox/ERT2^*mice with 4-hydroxytamoxifen (OHT) at indicated timepoints (E6.5-E10.5) and isolation of adult hematopoietic and tissue macrophage lineages followed by sample processing to retrieve *Polylox* barcodes. (**B, D, F, H, J**) Barcode heatmaps from *Rosa^Polylox/ERT2^*mice barcoded at E6.5-E10.5. *P_gen_*, barcode generation probability cutoff to select clonal barcodes; n = 2 mice per induction timepoint; barcode counts after P_gen_ cutoff/total barcodes detected on top of each heatmap. Individual barcodes in rows and sampled adult cell lineages in columns: mac., macrophages; Periton. Mac., Peritoneal macrophages; Mono/Granulo, Monocytes/Granulocytes; Hem. Prog., hematopoietic stem and progenitor cells. (**C, E, G, I, K**) Progenitor contributions (relative read counts) to sampled tissue macrophages (M, K, A, R) and hematopoietic progenitors (H) according to detected fate combinations. H: hematopoietic progenitors, M: microglia, K: Kupffer cells, A: alveolar macrophages, R: red pulp macrophages.

To focus on barcodes generated only in single progenitors (we refer to this as clonal barcodes because progeny from a barcoded progenitor constitute a clone, and all cells in this clone bear the same unique barcode), we filtered barcodes for low generation probability (*P_gen_*) as previously described (*15*). *P_gen_* cutoffs were adjusted for each induction timepoint to account for increasing growth, i.e. cell numbers, during embryonic development (see Materials and Methods: Post-sequence processing, Fig. S1C). At E6.5, we identified 555 clonal barcodes out of 598 total barcodes (92.8%) after *P_gen_* filtering in two mice across all sorted populations. Hence, we traced the fate of 555 single progenitors barcoded at E6.5, which gave rise to all analyzed tissue macrophages and hematopoietic lineages in the adult mouse.

In general, a high degree of barcode sharing between different cell types indicates a common progenitor, while barcodes unique for a single cell type identify independent origins. At E6.5, the by far most abundant barcode patterns were clonal barcodes that were shared across multiple cell populations, implying common, multi-lineage progenitors (Fig. 1B). We found that the vast majority of barcodes were reliably detected in sample repeats of the same population (Fig. S1D), hence barcodes were not undersampled. To quantify the lineage contributions of progenitors barcoded at E6.5, we focused on the most-studied macrophage lineages, i.e. microglia, Kupffer cells, alveolar macrophages, and red pulp macrophages, as well as on hematopoietic stem and progenitor cells (including Lin^−^Sca1^+^Kit^+^ cells, common myeloid progenitors, granulocyte-monocyte progenitors and megakaryocyte-erythrocyte progenitors). Using barcode read counts as proxy, we quantified the total contributions of distinct barcoded progenitors to adult populations (Table S1). The majority (>80%) of hematopoietic progenitors (H), microglia (M), Kupffer cells (K), alveolar macrophages (A) and red pulp macrophages (R) originated from a common multi- lineage progenitor (HMKAR) in E6.5 barcoded mice (Fig. 1C). In contrast, at this stage in development, fate-restricted progenitors, including those lacking microglia (HKAR progenitors), or generating microglia plus Kupffer cells (MK progenitors) made only minor contributions (Fig. 1C). In brief, common progenitors for all tissue macrophages and definitive hematopoiesis dominate during gastrulation (E6.5-E7.5).

### Lineage specification of tissue macrophages at the onset of organogenesis

Next, we asked when hematopoietic and tissue macrophage lineages diverge from one another in the embryo (*4*). To this end, we systematically introduced barcodes on consecutive days E7.5, E8.5, E9.5, or E10.5 (Fig. 1A). From adult mice, barcoded as embryos, macrophages and mature hematopoietic lineages were analyzed (Fig. 1D, F, H, J), and the barcode-based contributions of distinct progenitor fates were quantified (Fig. 1E, G, I, K). Progenitor barcoding was comprehensive with on average 75% (± 22%) of cells having a recombined *Polylox* barcode (Fig. S1B). Numbers of total clonal barcodes ranged between 348 and 1844 in the two mice for each timepoint (Fig. 1D, F, H, J). Barcode patterns and progenitor contributions were highly similar in each of the two barcoded mice analyzed for each induction timepoint, which demonstrates, next to robustness of the barcoding, temporal precision and consistent fate patterns underlying tissue macrophage developmental (Fig. S1E). Our experiments thus assessed the fates of several hundred progenitors per mouse in each time window.

The observed fate pattern and their eventual contributions to the different adult compartments depend entirely on the day of labeling as a ’viewpoint’ (Fig. 1B-K). While a common progenitor for all lineages prevailed on E6.5, subsequently, fate restrictions emerged. In particular, after E7.5 there was a sudden appearance of oligo- and unilineage fates. Prominent fate-restricted progenitors included a joint microglia and Kupffer cell (MK) progenitor. These bilineage progenitors contributed substantially more to adult microglia than to Kupffer cells. After E7.5, a common progenitor for Kupffer cells, alveolar macrophages, and red pulp macrophages (KAR) emerged. This KAR progenitor substantially contributed to these lineages at E8.5 and gradually diminished by E10.5. With developmental progression, a trace of macrophage potential was retained in progenitors for hematopoietic cells until E9.5 (e.g. HKAR, HKR) (Fig. 1G, I). Together, our data indicate a sequential fate restriction process, beginning with a multipotent progenitor at E6.5 and culminating in the emergence of unilineage progenitors independently generating hematopoietic progenitors, microglia, Kupffer cells, alveolar macrophages, and red pulp macrophages by E10.5 (Fig. 1K).

### Tissue-restricted residency of macrophages in fetal development

To trace macrophages after E10.5, and thus during fetal development, we induced barcodes on E10.5 and analyzed fetuses on E12.5, E13.5, and E14.5 (Fig. 2A, Fig. S4, S5). Therefore, we isolated F4/80^+^ macrophages from the fetal liver, and from cranial, trunk, and tail fragments (Fig. 2A, Fig. S4, S5). Macrophages from the liver and from these fragments showed predominantly unique sets of barcodes with minimal overlap (Fig. 2B, Fig. S5A, B), and, accordingly, low correlations with each other (Fig. 2C, Fig. S5C, D). Further linkage and read-based contribution analysis revealed stable uni-tissue barcoding for all examined compartments between E10.5 and E14.5, suggesting that most macrophages were labelled *in situ*, and were tissue-resident already in the fetus (see liver, cranial, trunk and tail fragment samples in Fig. 2C, D, Table S2). In contrast to macrophages, monocyte barcodes showed a high degree of linkage between monocytes from different tissues (see monocytes from the fetal liver, and cranial, trunk and tail fragments in Fig. 2C, Fig. S5C, D). Comparison of barcodes between fetal monocytes and macrophages revealed on average only 18% clonal barcode overlap and 25% read-based sharing (Fig. 2D, Fig. S5E, F). This barcode sharing may reflect a common origin of these cells after E10.5, or monocyte-derived macrophages. In any case, the majority of tissue macrophages were not fetal monocyte-derived, rather both were independent lineages. Further macrophage progenitor candidates, CD45^+^Kit^+^Lin^-^ progenitors with or without Csf1r expression (*4*, *9*), also lacked linkage to tissue macrophages (Fig. 2C, D), excluding a major role for these progenitors in macrophage generation after E10.5.

**Figure 2:**
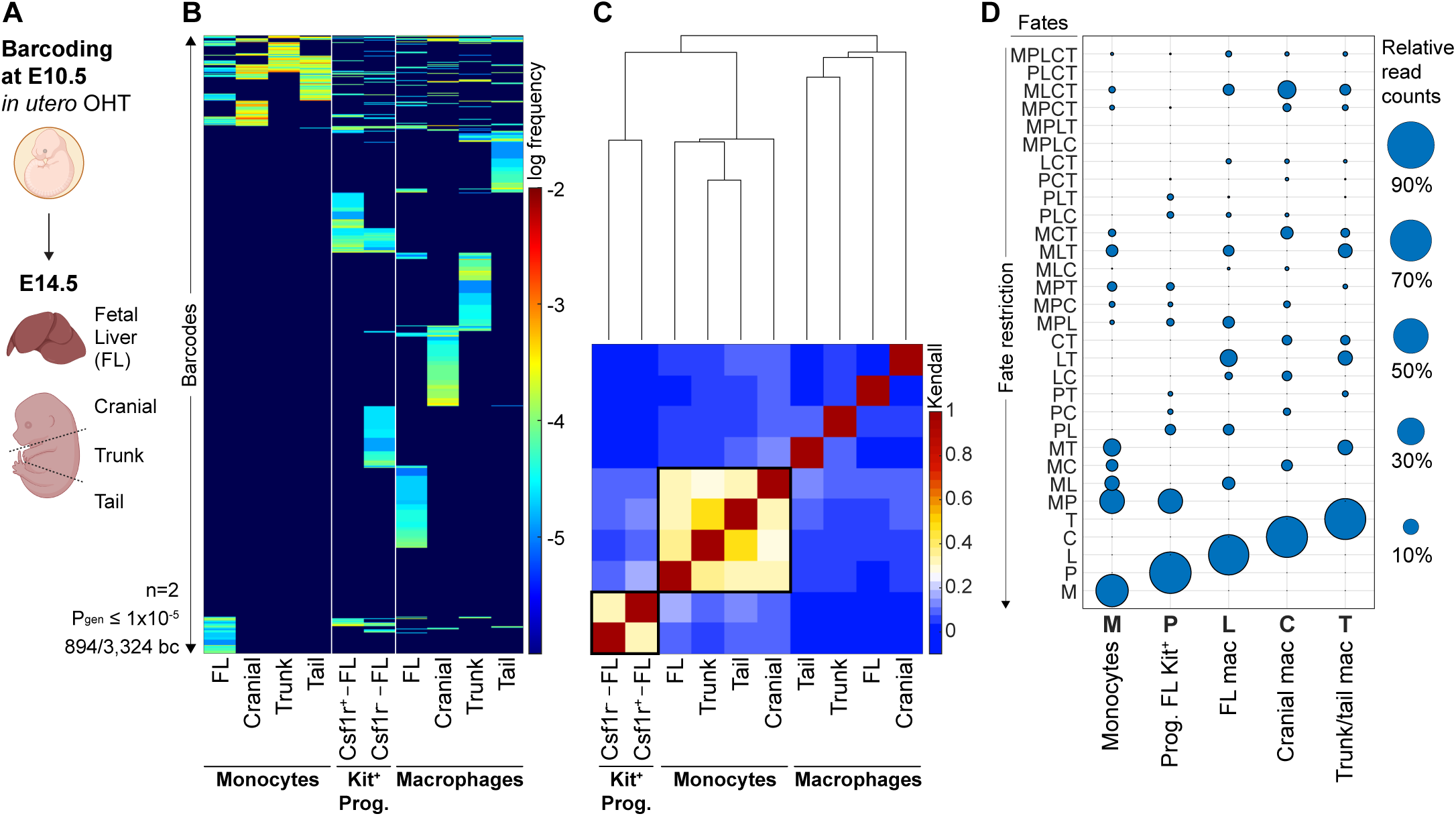
Correlation and quantification of barcode sharing of fetal monocytes and CD45^+^Lin^-^Kit^+^ progenitors with fetal macrophages. (**A**) In utero barcoding of *Rosa^Polylox/ERT2^* at E10.5 and analysis of different macroscopic fetal tissue regions at E14.5: Fetal liver (FL), cranial, trunk, and tail fragment of the embryo proper. (**B**) Barcode heatmap of E14.5 fetuses (n = 2) and comparisons of fetal monocytes, Kit^+^ progenitors (Prog.) from the fetal liver and macrophages from different fetal fragments. *P_gen_*, barcode generation probability cutoff to select clonal barcodes; barcode counts after *P_gen_* cutoff/total barcodes detected on the left bottom of the heatmap. Data displayed as (**C**) sample correlation heatmap using Kendall rank clustering of sampled populations and (**D**) bubble plot with contributions of progenitors with different fate combinations to the sampled populations. M = fetal monocytes (whole fetus), P = fetal liver Kit^+^ progenitors, L = fetal liver macrophages, C = cranial macrophages, T = macrophages from trunk and tail fragments.

### Inference of clonal differentiation pathways and dynamics

The distinct fates and contributions of progenitors to macrophage compartments across successive timepoints of embryonic development (Fig. 1B-K) suggest a reproducible, yet complex differentiation landscape. Using modeling based on our barcoding data (Fig. 1B-K), we define differentiation pathways from individual common macrophage (MKAR) progenitors by the lineage restriction events that occur in the expanding progeny of this progenitor (Fig. 3A, Fig. S6A). Principally, the several thousands of clones we measured (Fig. 1B-K) might develop according to one canonical differentiation pathway, or alternatively, macrophage progenitors could use several distinct lineage pathways.

**Figure 3.**
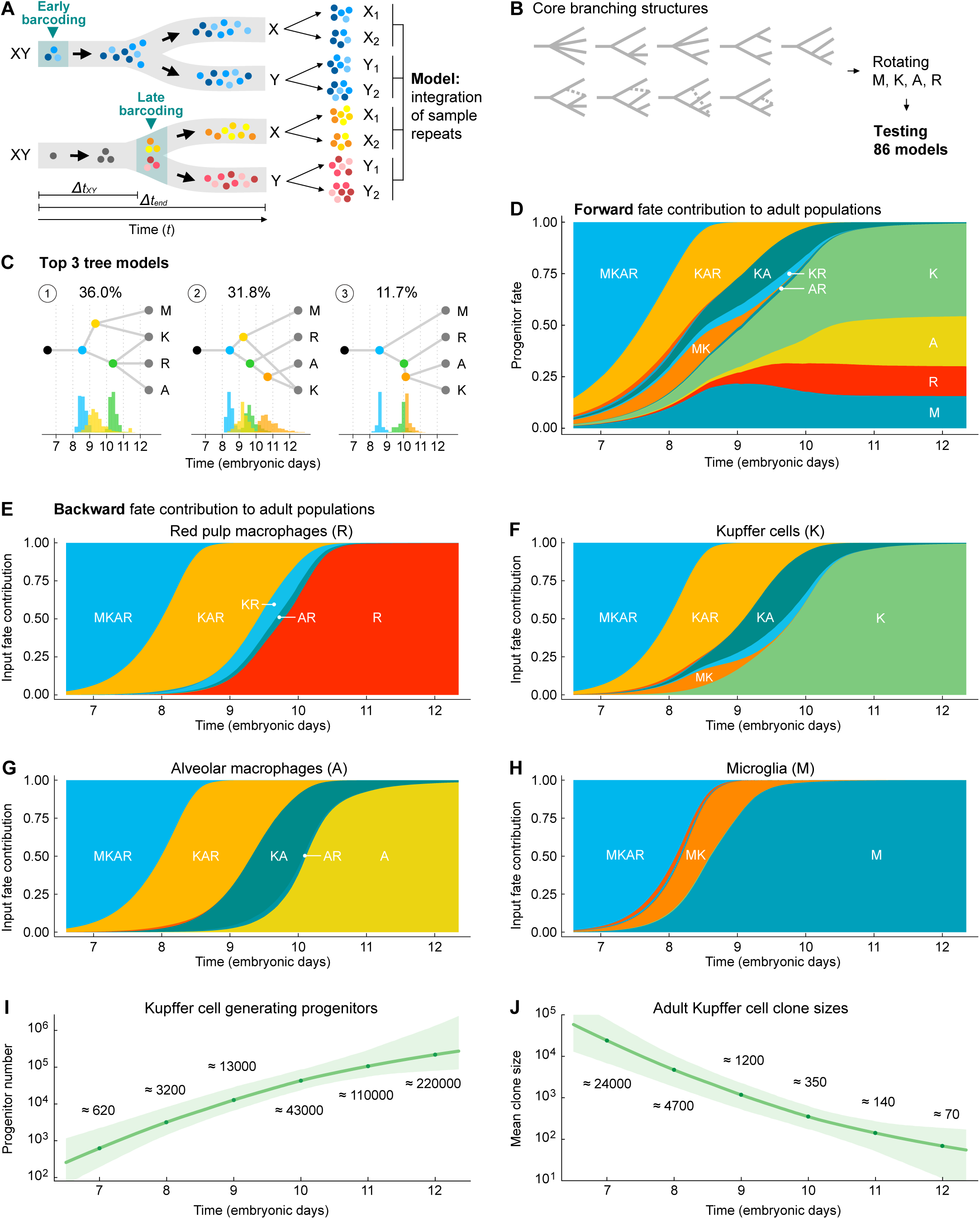
Computational inference of progenitor development and tissue macrophage lineage specification. (**A**) Framework of the mathematical model: Sample repeats (biological replicates of one population from an individual mouse) are compared for read-based barcode sharing to infer the clonal dynamics of lineage restriction. (**B**) Core branching structures. (**C**) Developmental programs of the top 3 models with inferred time windows of lineage restriction. (**D**) Integrated progenitor-centric fate map using the combined model average of all 86 models, the contribution of each model was weighted by its probability. (**E-H**) Retrospective population-centric fate map of backward progenitor contributions to red pulp macrophages (R), Kupffer cells (K), alveolar macrophages (A), and microglia (M). (**I**) Inferred progenitor numbers of Kupffer cells at different embryonic timepoints. (**J**) Inferred adult mean clone sizes of embryonic progenitors for adult Kupffer cells. (**I, J**) Mean Kupffer cell progenitor and clone size numbers for each timepoint are listed in Table S3.

Robust fate calling is a key condition for the classification of progenitor fates, and this is sensitive to progenitor sampling (see Materials and Methods: Mathematical modeling and inference). For example, fate combinations such as AR, KR, and KA (Fig. 1G, I, K) could correspond to clones originating from three distinct bilineage progenitors; alternatively, they could be incompletely sampled clones, with additional fates overlooked (e.g. a trilineage KAR progenitor). Hence, we controlled for sampling effects experimentally by repeat samples from every macrophage population (Fig. S1D, Fig. S6B). Moreover, in the case of the liver, we determined the absolute number of Kupffer cells, and the size of the samples drawn. We counted Kupffer cells in tissue sections of adult livers by immunofluorescent staining, microscopy, and image analysis (Fig. S7A-D), estimating a total of 1.5 × 10^7^ Kupffer cells. Hence, we analyzed 1.2% of the total adult Kupffer cell population for barcodes (Fig. S7E) which allows for robust barcode analysis (Fig. S8A).

To infer differentiation pathways from the experimental data, we devised a computational framework for simulating individual progenitor clones as they undergo proliferation and lineage restriction, factoring in experimental sampling to compare the *in silico* data with the experimental data (Fig. S6A). To visualize the principle, comparing information from early and late barcoding yields Δ*t*_XY_, the time at which a multilineage progenitor (X plus Y) gives rise to restricted (X or Y) progenitors (Fig. 3A, Fig. S6A). To fit the experimental data in an unbiased manner, we enumerated the differentiation pathways that could emerge from multi-fate MKAR progenitors, yielding 86 theoretically possible pathways (Fig. 3B). We then used Bayesian inference to fit the measured data of a total of 2444 individual clones (see Materials and Methods: Mathematical modeling and inference). The best model closely reproduced the experimentally observed, time-resolved barcode data (Fig. S6C). This model gave preference to a small number of related developmental pathways for the MKAR progenitor and fitted clonal expansion rates, thus making experimentally testable predictions.

Specifically, the model inferred the temporal succession of lineage-restricted progenitors, showing that only a small number of the theoretically possible fate combinations in progenitors are supported by the experimental data (Fig. 3C, Fig. S8B). Indeed, three tree models accounted together for 80% of the clones emanating from the MKAR progenitor (Fig. 3C, Fig. S8B). Accordingly, from MKAR progenitors, unilineage microglia (M) progenitors, bilineage microglia- Kupffer cell progenitors (MK) and trilineage progenitors for Kupffer cells, alveolar macrophages and red pulp macrophages (KAR) developed between E6.5 and E8.5 (Fig. 3D, Fig. S8C-E). To highlight one of these findings, the predominant bilineage MK progenitor pathway derived by the models (Fig. S8D) has not been reported previously and is clearly evident also in the barcoding data (Fig. 1E, G, I, K). After E8.5, the KAR progenitor gave rise directly to unilineage progenitors for Kupffer cells, alveolar macrophages or red pulp macrophages, or a bilineage KA progenitor (Fig. 3C, D). All other trilineage or bilineage progenitor populations were of negligible size, and, consistent with their paucity in the data, the model explains their occurrence by stochastic drop- outs of mature fates in small clones. We also took a mature-cell perspective and quantified which progenitors, over time, contributed to what extent to a specific mature macrophage population (Fig. 3E-H). In sum, our data and the modeling show a canonical differentiation pathway to tissue macrophages, composed of few, closely related tree structures.

The precise fraction for Kupffer cell sampling allowed us to infer the numbers of precursors that contribute to Kupffer cells as a function of time, increasing from ∼200 common macrophage progenitors at E6.5 to ∼70,000 Kupffer cell precursors by day 10.5 (Fig. 3I), when they could also be considered as premacrophages (*5*). The model further inferred that early progenitors (up to ∼E8.5) cycle about two to three times as rapidly as late progenitors (Fig. S9A, B). Unilineage Kupffer cell progenitors after E10.5 still undergo on average eight rounds of proliferative expansion (Fig. 3I, expansion from ∼70,000 to 1.5 × 10^7^ adult Kupffer cells, Fig. S9A). On this basis, the model predicts Kupffer cell clone sizes in the adult liver as a function of barcoding timepoints (Fig. 3J).

### Epitope barcoding with *Polytope* resolves spatial dynamics of Kupffer cell progenitor clones

To test Kupffer cell clone sizes experimentally, we employed a newly developed spatial fate-mapping system, called *Polytope* (*18*). The *Polytope* knockin locus (*Rosa26^Polytope^*) consists of nine distinct epitope tag cassettes flanked by loxP sites, allowing for random excision of cassettes upon Cre recombinase activity which is based on the principles of the *Polylox* system (Fig. 4A). The construct is expressed as an H2B fusion protein in the nucleus and epitope tags are detected by fluorescent immunostaining and multiplexed imaging. *Polytope* barcoding can generate up to 512 unique color codes, resolving spatial dynamics and sizes of clones in tissues.

**Figure 4:**
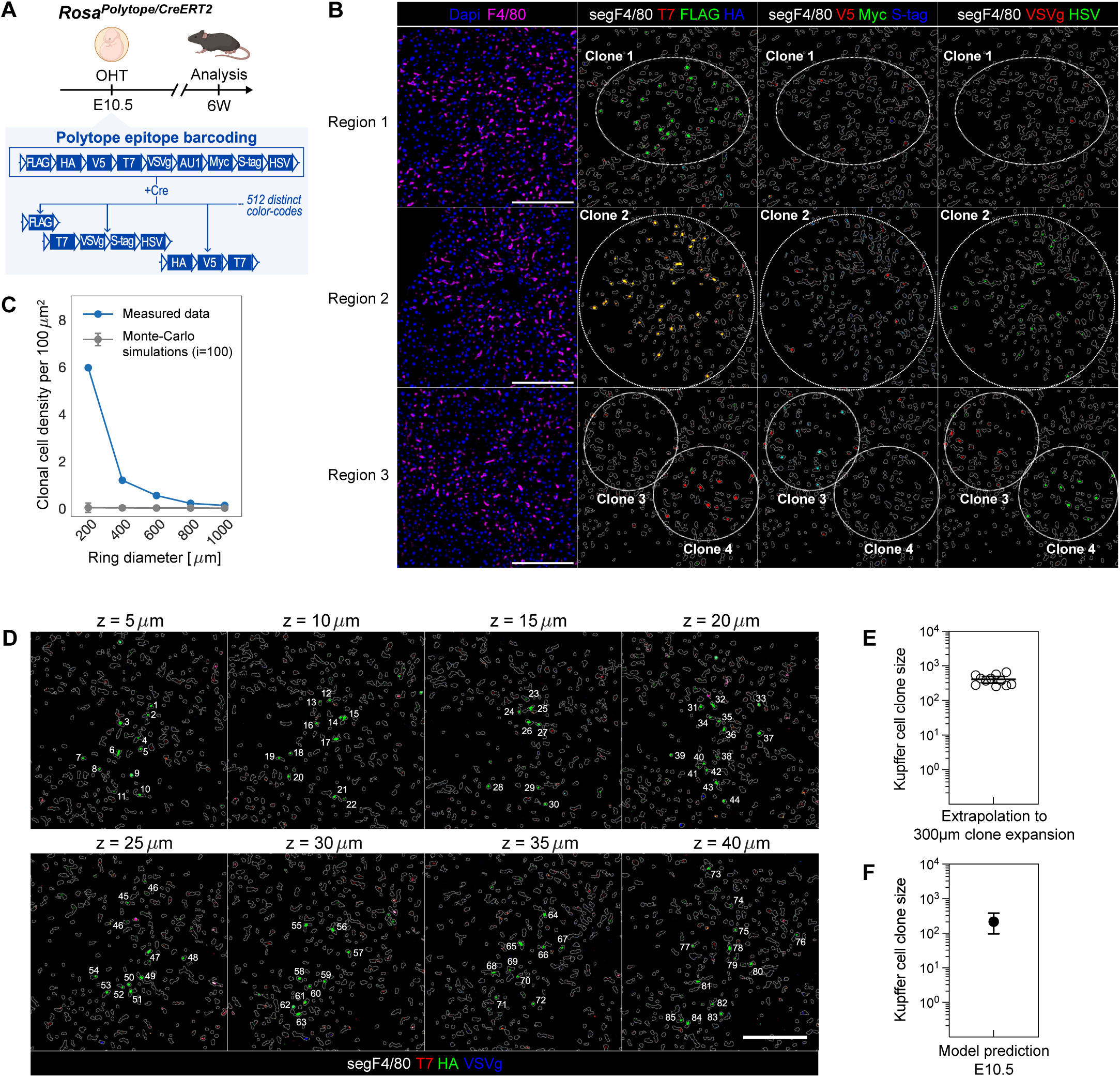
***Polytope* epitope barcoding for spatial fate-mapping of Kupffer cell progenitors from E10.5 to adulthood.** (**A**) *Polytope* color codes were ubiquitously induced *in utero* at E10.5 by OHT and analyzed in tissue sections of adult mice at 6 weeks of age. (**B**) Examples of color-coded Kupffer cell clones. Segmentation masks of F4/80^+^ Kupffer cells (segF4/80, white) indicate Kupffer cell boundaries. Depicted are only *Polytope* epitope barcodes within segF4/80 boundaries. Clone 1: FLAG^+^. Clone 2: FLAG^+^T7^+^HSV^+^. Clone 3: Myc^+^S-tag^+^VSVg^+^. Clone 4: T7^+^HSV^+^. Scale: 200 µm. (**C**) Clonal cell density of measured clones (n=11) and 100 Monte-Carlo simulations across concentric rings. Method adapted from Tay et al. 2017. (**D**) Consecutive sections for measuring Kupffer cell clone sizes (total z = 8 sections × 5 µm = 40 µm). Scale: 200 µm. (**E**) Extrapolation of measured Kupffer cell clone sizes detected with *Polytope* color codes in 40 µm (z-axis) to an average clone expansion of 300 µm. (**F**) Prediction of the mathematical model for Kupffer cell clone sizes at E10.5.

We ubiquitously induced color codes in *Rosa26^Polytope/ERT2^* embryos at E10.5, and analyzed liver cryosections at six weeks of age (Fig. 4A, Fig. S10A). Kupffer cell clones were locally restricted (examples shown in Fig. 4B). We measured the distance of each color-coded Kupffer cell to others of the same color code in two-dimensional sections, drawing concentric rings and counting the cells of the same color code (i.e., clone) within each ring (Fig. S10B). We compared these measurements to randomly distributed color codes, using one hundred Monte Carlo-simulated datasets to assess whether the observed clustering was due to chance (Fig. 4C). The measured data showed substantial enrichment of neighbor cells belonging to the same clone within 200 µm in diameter (Fig. 4C). Clone density decreased with distance but only approached random distribution at a 1000 µm radius, indicating that clones can span up to 800 µm (with most clones being 200-400 µm in size). We counted on average ∼400 cells per Kupffer cell clone (Fig. 4D, E, Fig. S10C), consistent with the model prediction (Fig. 4F). Using *Polytope* for spatial fate mapping, we hence arrived at a similar order of magnitude compared to the model-based calculations (Fig. 4E, F). In conclusion, epitope barcoding demonstrates the presence of Kupffer cell colony forming cells on E10.5, which expand locally, generating clones in the order of a few hundred cells in the adult liver.

### Spatial segregation of embryonic macrophage progenitors by E10.5

To test whether tissue locality is a general principle of early tissue macrophage development, we addressed this possibility by *Polylox* barcoding (Fig. 5). For barcode induction we chose one timepoint for shared barcodes (common progenitor at E7.5, see Fig. 1D, E), and one for unlinked barcodes (unilineage fates at E10.5, see Fig. 1J, K) between tissue macrophages. We analyzed adult mice by harvesting anatomically separated organ parts, including brain hemispheres, liver lobes, left and right lung, spleen halves, and left and right kidneys (Fig. 5A). In addition, we collected sample repeats from each tissue sample, except for the brain where we had one sample per hemisphere. As expected, after E7.5 induction, different samples from the same organ shared barcodes abundantly (Fig. 5B, C). In contrast, most barcodes were unique for organ fragments after E10.5 barcoding (Fig. 5B, C). To test whether the differences observed in E10.5 barcoded mice between organ fragments were ’true’ or reflected incomplete sampling, we analyzed the sample repeats from the same organ fragments (Fig. S11A). Indeed, these repeats showed substantially higher barcode sharing to each other than samples from different fragments (Fig. S11B, C). Collectively, these data are consistent with a common macrophage progenitor on E7.5 (Fig. 1D, E; Fig. 3D), tissue seeding by E10.5, followed by local expansion (Fig. 1J, K; Fig. 3D, Fig. 4). Consequently, once tissues are colonized, macrophages remain locally confined in their tissues.

**Figure 5:**
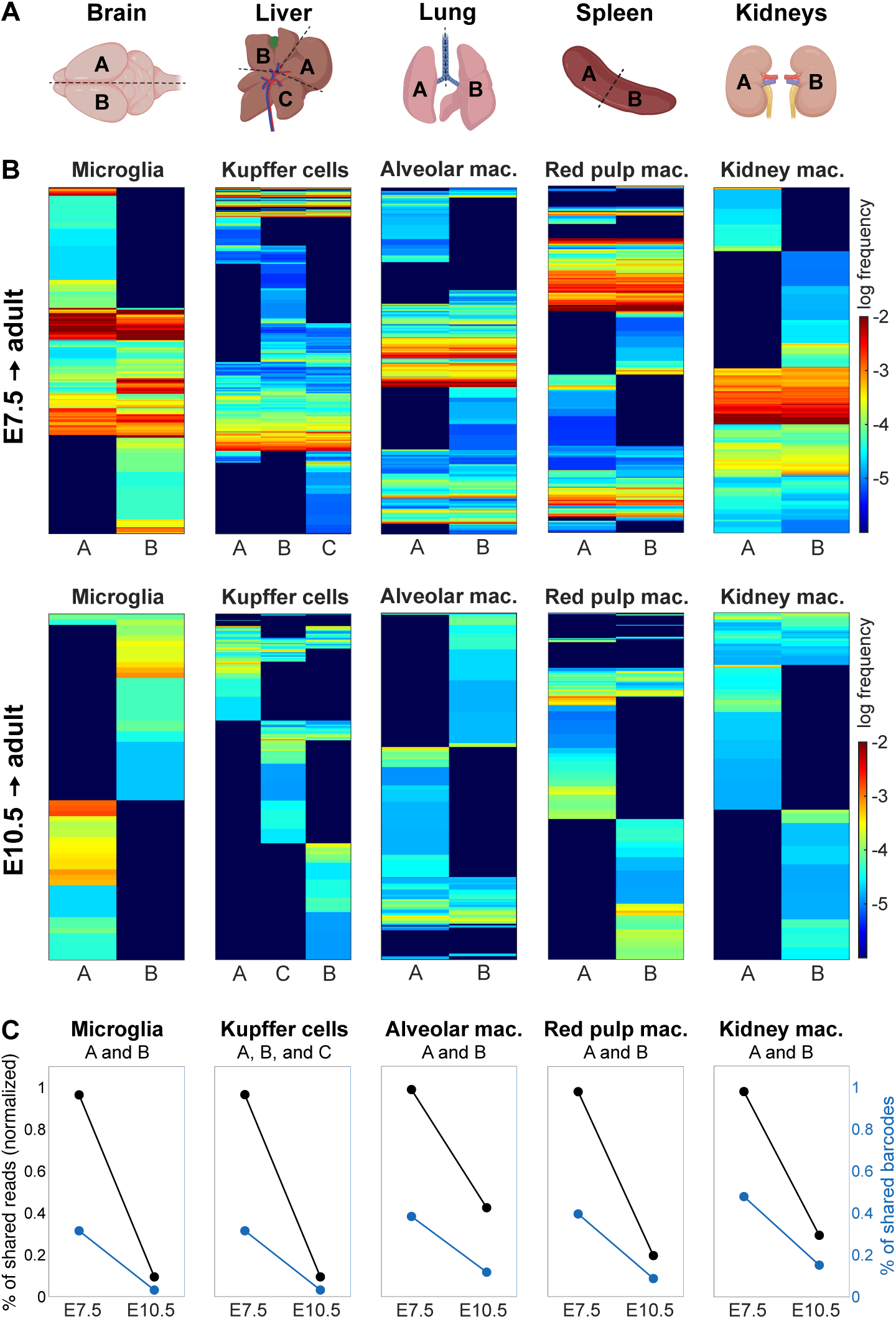
Local seeding dynamics by barcode analysis in organ fragments. (**A**) Analysis of organ fragments of brain hemispheres (A, B), liver lobes (A, B, C), left and right lung halves (A, B), spleen halves (A, B), and left and right kidney (A, B). (**B**) Barcode heatmaps of *Rosa^Polylox/CreERT2^* mice treated *in utero* with OHT at E7.5 (upper row) or E10.5 (lower row) and analyzed as adult mice. Comparison of microglia, Kupffer cells, alveolar macrophages, red pulp macrophages, and kidney macrophages of different organ fragments. Horizontal lines in the heatmaps represent individual barcodes; the color represents the barcode frequency in log-scale. Dark blue indicates that the barcode was not detected. *P_gen_* cutoff for E7.5 was ≤ 10^-3^ and ≤ 10^-5^for E10.5. (**C**) Quantification of barcode sharing as percent of shared reads to account for barcode frequency (left y-axis, black data points) and absolute number of shared barcodes (right y-axis, blue data points) at E7.5 and E10.5.

## Conclusion

Using holistic barcoding, we aimed at resolving the entire landscape of tissue macrophage development in the mouse. While their embryonic origin has been proven, the pathways leading to tissue macrophages in different organs remained controversial despite considerable experimental efforts. Barcoding across all developmental stages from gastrulation to organogenesis, and *a priori* independent of cell types or stages, now offers a powerful approach to comprehensively resolve pathways of macrophage development. Despite the fact that the initial progenitors are no longer present at the time of analysis, barcodes are retrospectively informative. In combination with mathematical modeling, we quantitatively and temporally mapped developmental trajectories of tissue macrophages between E6.5 and E10.5, and beyond. Barcode analysis revealed initially shared fates, followed by an array of fate combinations, and subsequently fate restrictions beginning on E8.5. Regarding microglia, our data are in line with their early development (*12*, *13*), however, in this pathway, we also find a novel progenitor with shared fate for Kupffer cells. The bulk of the other organ-specific macrophages have been proposed to be derived via fetal monocytes (*9*). Barcoding on E10.5, followed by analysis in fetuses between E12.5 and E14.5, revealed a clear developmental separation between tissue macrophages, fetal monocytes, and Kit^+^ progenitors. The infrequent barcode sharing between tissue macrophages and fetal monocytes may reflect a precursor-product relationship for few tissue macrophages, yet it may also result from a common origin of tissue macrophages and fetal liver monocytes. In any case, our data do not support the notion that fetal monocytes are the primary source of tissue macrophages. In addition to the lineage independence of macrophages and monocytes, the observed and modeled tree of macrophage development concludes its progression by E10.5, hence several days prior to the emergence of fetal monocytes. Barcode analysis in four regions of the fetus, and in fragments from adult organs demonstrates very early tissue residency and maintenance of a rather stationary life of tissue macrophages. Hence, neither in fetal nor adult mice appear tissue macrophages to migrate between organs or even within one organ. The measured fate patterns were highly reproducible between mice, suggesting intrinsic, tightly controlled developmental programs. This landscape provides a quantitative framework for investigations into the molecular determinants regulating fate decisions.

## Materials and Methods

### Mice

Homozygous *Rosa26^Polylox^* (*B6-Gt(ROSA)26Sor^tm1(Polylox)Hrr^*) and *Rosa^Polytope^* (*B6N-Gt(ROSA)26Sor^tm1(H2B-Polytope3.0)Jvr^*) were crossed to homozygous *Rosa26^CreERT2^* (*B6.129- _Gt(ROSA)26Sortm1(cre/ESR1)Tyj/J_*_) mice to generate *Rosa26*_*_Polylox/CreERT2_* _mice and *Rosa26*_*_Polytope/CreERT2_* mice, respectively. Lysed mouse tissue (Direct PCR-Lysis Reagent Tail, Peqlab) from ear stamps was used for genotyping by PCR. Mice were kept in individually ventilated cages under specific pathogen-free conditions in the internal animal facility at the DKFZ. Male and female mice were used equally, no randomization or blinding was used for experiments. All analyzed animals were included in the analysis except mice with a recombination of the *Polylox* casette below 10%. Sample size was not pre-determined. All animal experiments were performed in accordance with institutional and governmental regulations, and were approved by the Regierungspräsidium (Karlsruhe, Germany).

### Induction of *Polylox* cassette recombination

Timed matings of homozygous *Rosa26^ERT2^* and *Rosa26^Polylox^*were initiated in the evening and vaginal plug check was performed next morning and embryos of vaginal plug positive females were counted as E0.5. Pregnant females received a single dose of 4-hydroxytamoxifen (abbr. OHT, Sigma, H7904) combined with progesterone (Sigma-Aldrich) dissolved in peanut oil by oral gavage to induce *Polylox* barcode recombination. Doses varied between 0.3 mg at early and 1mg at later timepoints of embryonic development. Dosage of progesterone was always half of the OHT dosage. Treated mice were kept in separate ventilated cages and embryos were born by cesarean section at E19.5. Recombination efficiency was determined by PCR on 30,000 isolated blood cells after lysis of erythrocytes.

### Preparation of single-cell suspensions and fluorescence-activated cell sorting

Mice were euthanized by intraperitoneal injection of Ketamin (bela-pharm)/Xylazin (Elanco). We performed peritoneal lavage by washing the peritoneal cavity through a small incision of the abdominal wall with PBS and a pasteur pipette and transcardial perfusion using HBSS (Ca^2+^ and Mg^2+^ deficient) buffer containing Heparin (10 I.E./ml HBSS, ratiopharm). We isolated the brain, lung, liver, both kidneys, spleen and the whole skeleton for bone marrow isolation, and the following protocols were applied to prepare single-cell suspensions.

**Brain** was cut into small pieces and digested with papain from the ‘neural tissue dissociation kit’ (Miltenyi) for 15 min at 37 °C and an additional 15 min after adding the second enzyme mix. The tissue was homogenized by successively resuspending with a 10 ml serological pipette, 1 ml and 200 µl tips, and filtering with HBSS (containing Ca^2+^ and Mg^2+^) through a 70 µm cell strainer. Samples were centrifuged for 10 min at 300 × *g* and 4 °C and the resulting pellet was resuspended in 40% isotonic Percoll^®^ Cytiva™ (Sigma-Aldrich) and overlaid with HBSS (containing Ca^2+^ and Mg^2+^). Percoll gradient was centrifuged at 600 × *g* for 30 min without brake, and the pellet containing microglia was washed in HBSS.

**Lung, liver, and kidneys** were homogenized with scissors and digested in 10 mL RPMI including 1:10 Collagenase Type IV (60U f.c., Sigma-Aldrich), 25 µg/ml Dispase I (Roche), and 25 µg/ml DNase I (Sigma-Aldrich) for 45 min at 37 °C. Cell suspensions were filtered through a 100 µm cell strainer. The lung sample was lysed for 2 min in ACK (ammonium-chloride-potassium) lysis buffer to remove erythrocytes and washed with FACS buffer (PBS containing 5% fetal calf serum, Merck) before continuing with blocking and staining. Liver and kidney samples were centrifuged at 300 × g and 4 °C for 7 min, and the resulting pellet was resuspended in 10 mL HBSS. For the liver, a two-step Percoll gradient was prepared by underlying 25% Percoll solution with a 50% Percoll solution that was prepared with HBSS +/+ and centrifuged without brake and minimum acceleration at 1,800 × *g* and 4 °C for 20 min. Kidney cells were overlaid onto 20 ml of 30% isotonic Percoll solution and centrifuged at 400 × *g* and 4 °C for 3 min.

**Spleen** was cut into pieces and digested in 1 mL PBS containing 0.2 mg/ml collagenase D (Sigma-Aldrich), 25 µg/ml DNase I (Sigma-Aldrich) and 25 µg/mL Dispase I (Roche). After 20 min incubation in a heatblock at 37 °C and 800 rpm, samples were vortexed and resuspended with a P1000 pipette to dissociate cells. Dissociated single cells were transferred into 5 mL FACS Buffer containing 5 mM EDTA. Remaining sedimented tissue pieces were resuspended again in 1 mL digestion buffer and incubated again for 10 min before adding it to the single cell suspension. Samples were filtered through 70 µm cell strainer and passed over a pre-conditioned LS-column (using FACS buffer containing 2 mM EDTA) to isolate ferromagnetic red pulp macrophages. Column was washed with FACS buffer and then removed from the magnetic stand to collect isolated magnetic cells. Both separated samples were centrifuged at 1,200 rpm and 4 °C for 5 min and proceeded to blocking and antibody staining. The supernatant sample (depleted of magnetic red pulp macrophages) was splitted into two samples after Fc receptor blocking. A fraction was used for staining and subsequent sorting of T and B cells. To ensure efficient sorting of monocytes and granulocytes, the second sample was depleted of T and B cells by using magnetic Dynabeads (Life technologies). To deplete T and B cells, samples were stained for CD4, CD8, and CD19 for 30 min on ice and washed with PBS/5% FCS followed by depletion with magnetic Dynabeads (Life technologies) for 1 hour at 4 °C on a rotor according to manufacturer’s instructions. Cells that did not bind to magnetic Dynabeads were used for further blocking and staining and used to sort monocytes and granulocytes.

**Bone marrow** was isolated from the skeleton (femora, tibiae, fibulae, pelvis, humeri, radiuses, ulnae and spine) by crushing bones in FACS buffer using mortar and pestle and filtering the isolated cells through a 40 µm cell strainer. Fc receptors were blocked in PBS/5% FCS with 250 µg/ml purified IgG (Jackson ImmunoResearch Laboratories) for 20 min. Cells were stained for lineage markers (CD4, CD8a, CD11b, CD19, Gr-1, NK1.1, and Ter119) for 30 min on ice and washed with PBS/5% FCS before depletion with magnetic Dynabeads (Life Technologies) for 1 hour at 4 °C on a rotor according to manufacturer’s instructions. Cells that did not bind to magnetic Dynabeads were used for further blocking and staining.

**Embryos** were isolated in PBS on E12.5, E13.5, and E14.5 from the uterus. Embryos were decapitated to reduce blood in the circulation. Fetal livers were dissected, and embryos were separated into cranial, trunk, and tail/caudal fragments for further separate processing. Fetal livers were resuspended with a P1000 and filtered using a 100 µm cell strainer. The other three embryonic tissue regions were enzymatically dissociated for 20 min using 250 µg/ml Dispase I, 1 mg/ml Collagenase D, and 250 µg/ml DNase. Enzymatic reaction was stopped with PBS/5% FCS and centrifugation. Samples were resuspended in PBS/5% FCS and filtered for blocking and antibody staining.

Single-cell suspensions after digestion, depletion, or, in case of peritoneal cells, directly after lavage, were blocked in PBS/5% FCS with 250 µg purified IgG for at least 20 min. Cells were then stained with antibody cocktails in PBS/5% FCS at a concentration of 10^8^ cells/ml but in a minimum volume of 50 µl and for at least 30 min. Bone marrow stainings with anti-CD34 eflour 660 incubation were 90 min according to manufacturer’s instructions.

Cells were sorted with an LSR II Aria with a 100 µm nozzle and cells were collected in 300 µl PBS/20% FCS collection buffer. Samples were centrifuged twice for 5 min at 1,500 × g and turned by 180° in between. Pellets were resuspended in 25 µl lysis buffer containing 0.5 mg/ml proteinase K in nuclease-free water and PCR buffer.

### Definitions of isolated cell populations

All sorted populations were pre-gated for single live Lin^-^ cells. Lineage staining comprised for bone marrow CD4, CD8a, CD11b, CD19, Gr1, Ter119, and NK1.1. For liver, kidney, brain, RPM-magnetic enriched spleen, and peritoneum the lineage staining included CD3, CD19, Ter119 and for peritoneum additionally Siglec F. Splenic cells depleted of RPM were pre-gated for single live Ter119^-^ cells. Cell populations were defined as follows (Fig. S2, S3: Microglia (CD11b^hi^ CD45^lo^ Ly6C^-^), red pulp macrophages (CD45^+^ CD11b^lo^ F4/80^+^ MHC-II^+^ Tim4^+^), Kupffer cells (CD45^+^ CD11b^int^ MHC-II^+^ Tim4^+^), alveolar macrophages (CD45^+^ CD11b^lo^ F4/80^+^ CD11c^+^ Siglec F^+^), kidney macrophages (CD45^+^ CD11b^int^ F4/80^+^ MHC-II^+^), large peritoneal macrophages (CD45^+^ CD11b^hi^ F4/80^+^ Tim4^+^), monocytes in the liver (CD45^+^ CD11b^hi^ F4/80^-^ CD11c^-^ SSC^lo^ Ly6C^+^), granulocytes in the liver (CD45^+^ CD11b^hi^ F4/80^-^ CD11c^-^ Ly6G^+^), monocytes in the lung (CD45^+^ Siglec F^-^ MHC-II^-^ Ly6C^+^), granulocytes in the lung (CD45^+^ Siglec F^int^ Ly6G^+^). Splenic monocytes (CD11b^+^ CD115^+^ Ly6C^+^ CD19^-^ CD4^-^) and granulocytes (CD11b^hi^ CD19^-^ CD4^-^) were sorted from the sample after magnetic RPM isolation and T and B cell depletion. B cells (CD11b^-^ Ly6G^-^ CD19^+^), CD4 (CD11b^-^ Ly6G^-^ CD19^-^ CD8^-^ CD4^+^), CD8 (CD11b^-^ Ly6G^-^ CD19^-^ CD4^-^ CD8^+^) were sorted from splenic cells after magnetic RPM isolation and pregated. LSK (Sca-1^+^ Kit^+^) or LT-HSC (Sca-1^+^ Kit^+^ CD150^+^ CD48^-^), ST-HSC (Sca-1^+^ Kit^+^ CD150^-^ CD48^-^), and MPP (Sca-1^+^ Kit^+^ CD150^-^ CD48^+^) were sorted from lineage-depleted bone marrow cells, as well as MEP (Sca-1^-^ Kit^+^ CD16/32^-^ CD34^-^), CMP (Sca-1^-^ Kit^+^ CD16/32^low/int^ CD34^+^), and GMP (Sca-1^-^ Kit^+^ CD16/32^hi^ CD34^+^).

### *Polylox* barcode amplification and single-molecule real-time (SMRT) sequencing

Cells were processed as previously described (*19*). In short: Sorted cells were lysed with 25 µl lysis buffer (0.5 mg/ml proteinase K in nuclease-free water and PCR buffer) for 1 hour at 56 °C and 10 °C at 95 °C. Sample volumes were adjusted to 30 µl as input. The *Polylox* cassette was amplified by PCR using polymerase and buffer from the ‘expand long template PCR system’ (Roche), oligo no. 2427 (fwd primer, 5ʹ-CATACCTTAGAGAAAGCCTGTCGAG-3ʹ) and oligo no. 2450 (rev primer, 5ʹ-TGTGGTATGGCTGATTATGATCAG-3ʹ) and with the following program: 95 °C 5 min, 35x (95 °C 30 s, 56 °C 30 s, 72 °C 3 min), 72 °C 10 min. Samples were purified using AMPure XP beads (Beckman Coulter) according to the PacBio amplicon sequencing protocol and eluted in elution buffer (Qiagen). Library preparation for long-read sequencing on a Sequel machine (Pacific Biosciences) using barcoded adapters was performed according to the manufacturer’s instructions. Circular consensus sequence (CCS) reads were generated by the SMRT analysis software (Pacific Biosciences) and subjected to downstream processing.

### Post-sequence processing

Demultiplexed fastq files, containing CCS reads, were processed as previously described using the RPBPBR pipeline (*15*, *19*). In brief, the orientation of the barcode was determined by using the 5’ and 3’ end and trimmed, the bowtie2 (version 2.5.3) mapper was used for alignment of the nine forward and the nine reverse barcode segments and resulted in a .tsv file containing information on the number of total and intact barcodes as well as all detected barcodes and corresponding read counts. Using R, all samples from one mouse were then combined into one table containing barcodes (codes), read counts (reads), and sample names (annotation) and processed in MATLAB for barcode purging to filter for correct barcodes and to exclude all incorrect barcodes that violate the known modes of action of Cre recombination, e.g. barcodes containing multiples of one segment or having an uneven number of segment, also barcodes with unidentifiable segments were removed. Finally, the number of minimal recombination steps was assigned by a pre-calculated list, which in turn was then used to calculate the probability of generation (*P_gen_*) for each barcode. The *P_gen_* cutoff for barcode filtering was chosen based on reported total cell numbers in the embryo (Fig. S1C) and conservatively assuming (probably overestimating) that 10% of the total cells can be progenitors for tissue macrophages and hematopoietic cells. The final table containing barcodes, *P_gen_*, and read counts for each sample was used for downstream analysis for further calculations and to visualize the data in MATLAB or Python.

### Quantification of Kupffer cells

Livers of adult mice were perfused and fixed overnight in 4% paraformaldehyde, followed by dehydration in a 30% sucrose solution prepared in PBS. The volumes of eight fixed and dehydrated livers were measured using Archimedes’ principle with the 30% sucrose solution as a reference for later calculations. Liver pieces were then embedded in OCT Tissue-Tek (Sakura-Finetek) and stored at -80 °C. Livers were sectioned into 5 µm cryosections using a CryoStar NX50 and placed on Superfrost slides. The sections were blocked in blocking buffer (PBS/0.1% Triton X-100/1% BSA/10% donkey serum, Abcam, ab7475) for 1 hour at room temperature in a humid chamber. This was followed by incubation with a rat anti-F4/80 antibody (1:100, Invitrogen, BM8, MF48000) in staining buffer (PBS/0.1% Triton X-100/1% BSA) overnight at 4 °C. The samples were washed four times with staining buffer before applying the secondary antibody (1:800 donkey anti-rat IgG H&L Alexa Fluor-647, Abcam, ab150155) for 1 hour at room temperature. This was followed by DAPI counterstaining, PBS washing, and mounting with Fluoromount-G (Southern Biotech, 0100-01). Liver sections were imaged using a Keyence BZ-X810 fluorescence wide-field microscope equipped with a 20X plan apochromate objective (Keyence, BZ-PA20). Images were stitched using the integrated software. Image analysis was performed in ImageJ, including preprocessing steps such as background subtraction and mean filtering. Nuclear segmentation was conducted on the DAPI channel using global Li thresholding, and macrophage segmentation was done on the F4/80 channel using the same method. The number of F4/80^+^ cells was determined by counting the segmented nuclei that showed a 99% pixel overlap with the F4/80 segmentation mask. The total area of the imaged tissue was measured by segmentation on contrast-enhanced images (autofluorescent background).

### Mathematical modeling and inference Code and method availability

All code was written in the Julia programming language and is available on Github. Methods for generative models with four terminal populations are contained in the package LineageBranching.jl and executing code for the macrophage inference is contained in LineageBranchingMacrophages. The Bayesian inference was conducted with ABCdeZ.jl. https://github.com/hoefer-lab/LineageBranching.jl https://github.com/hoefer-lab/LineageBranchingMacrophages https://github.com/hoefer-lab/ABCdeZ.jl

### Inferring clonal dynamics from in situ barcoding data: a single lineage

The measured barcoding patterns are generated by the dynamics of macrophage progenitor cell proliferation and differentiation. We model these dynamics based on the theory of stochastic branching processes and use approximate Bayesian computation for model selection and parameter estimation from the experimental data.

We begin by considering a single mature cell population that is composed of an ensemble of clones (measured via barcodes) and ask how many barcodes will overlap in two sample repeats; the mathematical result will establish a relation between measurable barcode overlap and the experimentally unknown expansion of the clones. Specifically, we assume that the clones, each originating from a single progenitor cell, have grown with an average cell division rate !. We now show that measuring the barcode overlap and sampling frequency yields an estimate of the mean number of cell generations, !", that the clones have undergone in the time interval " between progenitor barcoding and sampling of mature progeny. For simplicity, we neglect the likely very small death rate of dividing progenitors in the fetus and model clonal expansion as a pure birth process with a mean generation number λt. A clone will reach the final size *X_t_* = *x* with probability *p*(*x*) = (1 − *b*) *b*^"*x*-1^, where *b* = 1 − exp(−λt), and x = 1, 2, 3, … (*20*). Accordingly, the mean clone size is E(*X_t_*) = exp(λt) cells. Based on these statistics, we now compute the expected overlap of barcodes in two small samples from a mature cell population. In both samples together, we recover 5 distinct clones (i.e., clonal barcodes) and define the sample overlap as

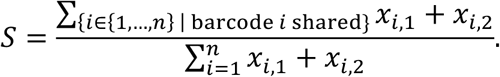

This expression jointly accounts for sharing and clone size, as *x_i,j_* is the sampled clone size of barcode *i* in sample repeat j ∈ {1, 2}. Specifically, each sample *j* consists of a fraction *p*_j_ of cells from the total population, such that *p*_1_ + *p*_2_ + q_loss_ = 1 , where q_loss_ is the probability to not sample a cell. For a sufficiently large number of clones (practically, 10 clones or more), applying the law of large numbers and the continuous mapping theorem yields the following analytical relation between the expected sharing, the mean generation number, and the sampling fractions

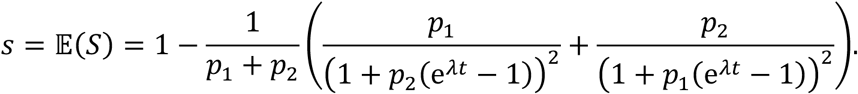

For sample repeats of equal size (*p*_1_ = *p*_2_), we find

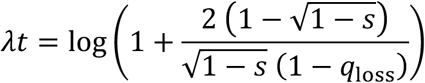

which relates progenitor expansion to sharing of sample repeats and experimental sampling loss. Hence, measuring of *s* (as done in all our barcode analyses) and *q_loss_* (as done for Kupffer cells) yields the mean number of cell generations from the progenitors to the mature cell populations (see Fig. S9A). In turn, this predicts Kupffer cells clone sizes of in the adult liver (Fig. 3J), which we confirmed experimentally (Fig. 4).

### Generative modeling of macrophage development

Having quantified barcode sharing in sample repeats of the same population, we now turn to multiple macrophage lineages. The magnitude of barcode sharing in samples across different lineages will inform about lineage restriction events in differentiation pathways of macrophage progenitors. In particular, less sharing across lineages than across sample repeats will indicate a lineage split between the times of barcode induction and measurement, as detailed in the following.

We systematically developed generative models of fate restriction during the developmental expansion of macrophage progenitor clones. Each model is specified by three types of parameters: (i) the kind and order of lineage restriction events, (ii) the timing of lineage restriction events, and (iii) clonal expansion.

### Division phases and differentiation events in the generative models

We modeled the progenitor differentiation pathways starting from a common progenitor population that expands with a given rate λ until the first differentiation (i.e., lineage restriction) event. The differentiation events are modeled as distributing the progenitor cells, at differentiation time *t*’ into two or more lineage-restricted progenitor populations. Once distributed, the clones in each progenitor population expand until the next differentiation event. This process is repeated until all unilineage macrophage progenitors are specified. The expansion of clones between lineage restriction events was modeled as a pure birth process by drawing clone sizes from a shifted negative binomial distribution with mean *x*_0 <_ exp(λΔt) on integer support {x_0_, x_1_ + 1, x_0_ + 2, … }. This is much more time-efficient than stochastic simulation. The fate restriction events, that distribute the cells from the current progenitor population (e.g., with fate AB) through a split into the two subsequent populations (e.g., with fates A and B, respectively), were realized as a binary sampling event for each progenitor cell with defined split probabilities (e.g., *P_A_* and 1 − *P_A_*, leading to a binomial distribution). Triple and quadruple lineage splits were realized as immediate successive binary splits. Given a certain lineage topology, this process of division phases followed by lineage splits was repeated until the final clone sizes, before experimental sampling, of all terminal populations were obtained. For topologies that have different progenitor routes leading to the same terminal population, the routes were simulated separately with the same iterative procedure, and the clones of a fate with multiple input routes were merged after the last division phase before the experimental capture (as the pure birth process is a branching process of statistically independent clonal expansions, the exact time of merging in the final phase is irrelevant). To compare simulations with experimental data, we drew sample repeats for each population. For this, the terminal clone sizes were subjected to multinomial sampling into three categories for sample repeat 1, sample repeat 2 and a third category for not-sampled / lost cells, with associated sampling probabilities *P*_1_ = *P*_2_ = (1 − *q*_loss_)/2 (symmetric sample repeats) and p_3_ = q_loss_. We used these generative models to simulate the lineage restriction process from a set of common progenitor cells (with x_0_ ≥ 1) to the final adult macrophage populations.

### Barcode introduction in the generative models

To address the *Polylox* barcoding data, focusing on clonal barcodes (i.e., the progeny of single, uniquely barcoded cells), we computationally introduced clonal barcodes, with x_0_ = 1, at times defined by the experiments. Specifically, we simulated each of the independent experiments (10 in total, 2 mice per induction timepoint, for induction times E6.5, E7.5, E8.5, E9.5, and E10.5). A progenitor population without barcodes was simulated until the induction time (*t*_ind_) of the given experiment. At these times a number (n*t*_ind)_ of clonal barcodes was introduced within the simulated total population. To account for the time window of barcode recombination by OHT-activated Cre (*17*, *21*), we took "_?@4_ to be 12 hours after the timepoint of OHT application (i.e., E7, E8, E9, E10 and E11). A clone was established by specifically marking a single cell for the subsequent simulation and removing it from the remaining unrecombined cells (the previous total). The simulation was completed by iterating the remaining phases of divisions and lineage restriction, and tracking barcode identities. If multiple progenitor populations existed during the time of barcode induction, the barcodes were introduced with uniform probability in all the existent cells across the populations. The simulation result, per experiment, is a collection of simulated clone sizes for an array of clonal barcodes, with two sample repeats for each terminal lineage.

### Parameters in the generative models

The differentiation pathway topology is fixed for a given model; we enumerate all possible topologies and systematically select models accounting for the experimental data (see the next section). For each model topology, we specify the following set of parameters by Approximate Bayesian computation using the experimental data: 1) the times of lineage restriction events (e.g., Δ*t_AB_* for an “AB to A and B” model), 2) the total time span (Δ*t_end_*) from the emergence of the common progenitors to establishing all mature tissue-resident macrophage populations, 3) the split probabilities into the different lineages for each restriction event (e.g., *p*_A_ for an AB progenitor to give rise to lineage A; note that in this example *P_A_* + *P_B_* = 1), 4) the division rates for each progenitor and terminal population, and 5) the number of introduced clonal barcodes (Δ*t_end_*) ). In detail, two division rates were introduced, λ_pre_ and λ_terminal_, respectively for multi- or oligopotent progenitors and terminal lineage expansion. This allows for time dependence of division rates through development at a resolution that can be inferred from the experimental data. The number of initial cells in the common progenitor population at E6.5 was set to the maximum of the clonal (“rare”) barcode number parameters (n_tind_ for each induction time t_ind_) to guarantee that as many clonal barcodes can be introduced in the simulation as was seen in the respective experiment. The sampling parameter q_loss_ was set to 99.6% corresponding to a coverage per sample repeat of 0.2% for each terminal lineage (30,000 cells sorted per sample out of ∼15 million total adult cells, as experimentally determined for Kupffer cells, Fig. S7); in separate computations, we also studied the effect of parameter q_loss_by systematically varying it (Fig. S8A).

### Mean progenitor clone sizes from the generative models

The thus specified model simulations realize sequential stochastic birth processes with sampling at the final timepoint. In topologies in which each population has precisely one type of progenitor, the clone size distributions are shifted geometric distributions on support {1, 2, 3, … }, for surviving clones, mixed with a delta distribution at zero clone size, for clones that have gone extinct in the process. Analytical expressions for the mean clone sizes for the terminal populations can be computed by iterating through the individual phases. For example, in an “AB to A and B” lineage branching process, the mean for terminal lineage “A” is

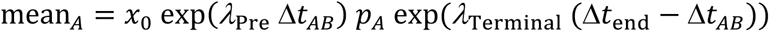

(before experimental sampling). If a population has more than one type of progenitor, the mean clone size is obtained by adding the mean clone sizes of the individual routes feeding this population.

### Creation of model set

To probe a diverse set of potential lineage restriction processes, a set of nine core topologies was devised that define distinct routes by which a common progenitor population eventually gives rise to four terminal populations (Fig. 3B; core topologies 1 to 5 in upper row, 6 to 9 in lower row, from left to right). The terminal lineage branches were then associated with the labels for the macrophage populations (M, K, A, R) in every possible combination, yielding a total of 86 unique topologies. The number of models per core topology is as follows: c1=1, c2=4, c3=6, c4=3, c5=12, c6=12, c7=12, c8=24 and c9=12.

The core topologies include all strictly tree-like topologies in which each product population has exactly one progenitor type. We allowed for binary, as well as triple or quadruple lineage splits. Further core topologies were generated by allowing for an additional path to an already existing terminal lineage, under the constraint that the added path does not lead to duplicate up- or down-stream progenitor fates in the resulting topology. Thus, core topologies c6 to c9 were obtained.

### Statistics for multi-lineage inference

As the basis for the inference, we derived statistics from the barcoding data of ten embryonically induced mice, two mice per five induction times (E6.5, E7.5, …, E10.5), as follows. Only clonal barcodes were used. For each mouse, a pair of (biological) sample repeats (from the same location) for each terminal lineage was selected, comprising up to eight sample columns (M1, M2, K1, K2, A1, A2, R1, R2); technical repeats (resequencing) were merged beforehand if available. For a given mouse, a primary statistic was constructed by quantifying the read-based normalized progenitor contributions to the eight sample columns according to their detected fate combination (255 fates, i.e., 2^8^-1). In the few cases where sample repeats were not available for a terminal population, the statistic was reduced to six or seven sample columns and 63 or 127 fates, respectively (e.g., M only instead of M1 and M2; the generative model was adapted accordingly in these cases). A secondary statistic contained the number of sampled clonal barcodes, total over all selected samples per mouse, to enable the inference of the original clonal barcode number introduced in vivo (nt_end_ ).

### Inference over model set

A Bayesian framework was employed to infer parameters and model probabilities, updating prior assumptions with the experimental data to obtain posterior knowledge. A simulation-based, like-lihood-free inference procedure was used. Tracking prior particle weights within common ABC-SMC (Approximate Bayesian computation, Sequential Monte Carlo) algorithms allowed to obtain model evidence estimates for model comparison, in addition to posterior parameter samples. This inference code was packaged into ABCdeZ.jl, and applied to each model within the model set (86 models total). For each model, a posterior model probability and Bayes factor were obtained based on its estimated evidence, to compare it within the overall model set.

A discrete-uniform model prior was chosen, and within each model, the parameter priors were uniform with the following support: a) continuous division rates within 0 and 3/day, b) continuous time spans for all birth phases and the total time span (Δt_end_) within 0 and 30 days, c) continuous split probabilities within 0 and 1, and d) the discrete number of clonal barcodes introduced in vivo per induction time (n_tend_) between 1 and 2000. The prior was further pruned to eliminate impossible parameter combinations (specifically that the total time span could accommodate all barcode induction and lineage restriction times), and to limit the mean clone size of a single E6.5 progenitor to no more than 10^8^ cells total.

To enable likelihood-free inference by ABCdeZ.jl, the distance of the data statistics (the same for all inference runs and iterations) and the model statistics (repeatedly generated) were summarized by a single value ≥ 0 in the inference iterations. To keep rich distribution information, data and model statistics were compared by the Jensen-Shannon divergence (a symmetrized and smooth-ened Kullback-Leibler divergence) for all ten experiments; divergences were then normalized by the sum of the respective data statistics, and the mean of all normalized divergences was reported as a single distance value.

### Posterior analyses

Re-simulations were subsequently used to generate posterior insights, drawing model and parameters according to the inferred posterior model probabilities and, within each model, posterior parameter distributions. If not otherwise stated, such Bayesian model-averaged posterior knowledge is supported by the total model set. Most importantly, they provide insights into the lineage restriction process in vivo, undoing the experimental sampling effect of having analyzed only a fraction of the total adult tissue populations.

#### Fate maps

Forward progenitor fate maps were generated by introducing single-cell progenitors continuously over a time window from ∼E6.5 to ∼E12.5 in posterior simulations and then tracing and analyzing their final fate in the adult tissues before experimental sampling. Backward progenitor fate maps were computed analogously by quantifying the final composition of each terminal tissue in terms of the cell-based fractions of all incoming progenitor fates.

#### Clone sizes

Adult mean clone sizes of embryonic progenitors were again based on the continuous introduction of single-cell progenitors in posterior simulations, and then computed as mean values of the simulated clone sizes in the terminal populations before experimental sampling (with or without conditioning on lineage specificity, depending on context). Predictions of Kupffer cell in the embryo were obtained by relating the computed mean clone sizes of Kupffer cell progenitors with the measured ∼15 million total Kupffer cells in the adult liver.

#### Division rates

Posterior distributions of division rates were obtained by aggregating λ_pre_ and λ_terminal_ parameters from the posterior re-simulations. Note that, as an orthogonal approach, (early and late) division rate estimates were also computed from sample repeats only, across all terminal populations and mice (see section above ‘Inferring clonal dynamics from in situ barcoding data’, and Fig. S9A).

#### Goodness of fit

To inspect the goodness of the fit, the primary statistics of the progenitor fate contributions (“bubble charts”) were computed for posterior re-simulations with the same barcode induction times and experimental sampling parameters as in the inference and averaged over the simulations for model-averaged statistics per induction timepoint.

#### Developmental programs

Depictions of developmental programs were based on the inference results for models individually, with the inferred model probability, the employed model topology, and the inferred posterior lineage split times as indicated (Fig. 3C and Fig. S8E).

#### Effect of sampling

We performed robustness analysis with respect to experimental sampling by repeating the inference with the top-ranked model for varying experimental sampling rates &_$_ and &_’_ both set to 0.2% (regular), 0.002% or 20%, and computing the posterior progenitor fate map for this single model only, otherwise as described above (Fig. S8A). Noteworthy, varying sampling rates show the overall robustness of progenitor fate maps.

### Multiplexed imaging of liver cryosections from induced *Polytope* mice

At six weeks of age, mice were euthanized and perfused with PBS through the left ventricle, before organs were collected and fresh frozen in OCT using liquid nitrogen chilled 2-methylbutane with subsequent storage at -80°C. Blocks were cut as 5 μm sections and placed on SuperFrost Ultra Plus GOLD glass slides (Epredia) before being stored at -80°C after 1h drying at room temperature. Before immuno-staining, sections were dried for 30 min at room temperature, followed by 15 min fixation with 4% PFA/PBS and three consecutive washing steps in PBS, and subsequent solubilization in PBST (PBS/0.1% Triton X-100) for 10 min at room temperature and blocking in blocking buffer (PBST/1% BSA/10% donkey serum) for 1h at room temperature. A complete list of the antibody panel is provided in Table 2 and multiplexed imaging was performed as described in the manual protocol of IBEX (*22*). Briefly, directly conjugated antibodies (Alexa Fluor 488 or 555 or 647), were incubated at 4°C overnight in staining buffer (PBST/1% BSA) with subsequent washing with staining buffer, PBST, incubation with 1 μg/mL Dapi/PBST and washing with PBS, before mounting in Fluoromount-G/PBS (50%/50%). Whole section images were acquired on a Zeiss Axioscan 7 (20X objective/ 0.8 plan-apochromat air). Coverslips were removed by demounting in PBS, sections were washed in PBS and photobleached in 1.0 mg/mL LiBH_4_ solution (dissolved in diH_2_O) for 20 min at room temperature under exposure to a white LED light (daylight lamp). Samples were washed three times in PBS and incubated in blocking buffer for 30 min at room temperature before the next IBEX cycle started. Effective bleaching was confirmed on a widefield fluorescent microscope (Keyence, BZ-X810).

**Table 1:** Antibodies for fluorescence-activated cell sorting.

| Antigen | Clone | Fluorophore | Vendor | Dilution |
| --- | --- | --- | --- | --- |
| CD3 | 17A2 | BV421 | BioLegend | 1:200 |
| CD4 | GK1.5 | BV421 | BioLegend | 1:400 |
| CD8a | 53-6.7 | BV421 | BioLegend | 1:400 |
| CD8a | 53-6.7 | PE | BD Pharmingen | 1:200 |
| CD11b | M1/70 | BV421 | BioLegend | 1:400 |
| CD11b | M1/70 | PE-Cy7 | eBioscience | 1:400 |
| CD16/32 | 93 | PE-Cy7 | eBioscience | 1:400 |
| CD19 | 1D3 | APC | BD Pharmingen | 1:400 |
| CD19 | 6D5 | BV421 | BioLegend | 1:200 |
| CD45 | 30-F11 | A700 | BioLegend | 1:100 |
| CD48 | HM 48-1 | A700 | eBioscience | 1:400 |
| CD115 (Csf1r) | AFS98 | BV605 | BioLegend | 1:400 |
| CD150 | TC15-12F12.2 | PE | BioLegend | 1:200 |
| F4/80 | BM8 | BV711 | BioLegend | 1:100 |
| Gr-1 | RB6-8C5 | BV421 | BioLegend | 1:400 |
| CD117 (Kit) | 2B8 | BV711 | BioLegend | 1:800 |
| Ly-6C | HK1.4 | APC/Fire™750 | BioLegend | 1:200 |
| Ly-6G | 1A8 | PerCP-Cy<sup>TM</sup>5.5 | BD Pharmingen | 1:100 |
| MHC-II | M5/114.15.2 | APC | eBioscience | 1:400 |
| NK1.1 | PK136 | BV421 | BioLegend | 1:200 |
| Sca-1 | D7 | PerCP-Cy5.5 | eBioscience | 1:200 |
| Siglec F | E50-2440 | BV421 | BD Pharmingen | 1:100 |
| Siglec F | E50-2440 | PE | BD Pharmingen | 1:100 |
| Ter119 | TER-119 | BV421 | BioLegend | 1:200 |
| Ter119 | TER-119 | FITC | eBioscience | 1:100 |
| Tim4 | RMT4-54 | PE | BD Pharmingen | 1:200 |

**Table 2:** Antibodies for multiplexed imaging with the IBEX protocol.

| IBEX cycle | Antigen | Clone | Fluorophore | Vendor | Dilution |
| --- | --- | --- | --- | --- | --- |
| 1 | AU1 | AU1 | AF488 | Biolegend (901901) | 1:400 |
|  | VSVg | E8S5G | AF555 | Cell Signaling Technology | 1:600 |
|  | T7 | D9E1X | AF647 | Cell Signaling Technology | 1:600 |
| 2 | HSV | polyclonal | AF488 | Novus Biologicals (NB600-513-AF488) | 1:400 |
|  | S-tag | D2K2V | AF555 | Cell Signaling Technology | 1:800 |
|  | Myc | 9B11 | AF647 | Cell Signaling Technology | 1:800 |
| 3 | V5 | SV5PK | AF488 | abcam (ab206566) | 1:250 |
|  | HA | C29F4 | AF555 | Cell Signaling Technology | 1:600 |
|  | FLAG | FG4R | DyL650 | Invitrogen (MA1-91878-D650) | 1:800 |
| 4 | F4/80 | BM8 | AF647 | Biolegend (123122) | 1:100 |

### Image analysis of IBEX image data

Raw images were preprocessed by background subtraction (median filtering) and images from IBEX cycles were aligned using the MultiStackReg plugin in ImageJ based on the Dapi channels (https://github.com/miura/MultiStackRegistration.git). Two segmentation masks were created: A nuclear segmentation mask using Cellpose (pre-trained generalist model for nucleus segmentation v1.0) on the Dapi channel, and a Kupffer cell segmentation mask based on the F4/80 channel using global thresholding (*23*). The nuclear segmentation mask was used to measure mean fluorescent intensity values for all cells. Segmented nuclei overlapping ≥99% pixel area with the Kupffer cell segmentation mask were considered Kupffer cell nuclei. *Polytope* color codes were retrieved from the mean fluorescent intensity dataset as described in (*18*) using a density-based spatial clustering of applications with noise (DB-SCAN) approach to detect cells with similar epitope tag mean fluorescent intensities and their corresponding (x, y) coordinates. For testing local clonal expansion, the dataset was analyzed by mapping the distribution of cells by drawing concentric rings around a central point (100 μm increments up to 1200 μm) and counting the number of clonal cells (same color-code) within each ring (Fig. S10B). Counts were normalized to the corresponding area (μm^2^) of the ring (*24*). In order to distinguish between random and clonal expansion based on their spatial organization, we performed 100 Monte Carlo simulations by randomly distributing detected color codes over the Kupffer cell (x, y) space and repeating the measurements.

## Acknowledgements

We thank all members of the Rodewald and Höfer group for ongoing support and discussions, Sven Schäfer for technical support, and the team of animal caretakers at DKFZ for support and expert animal husbandry.

## Funding

L.F. was supported by the I&I Helmholtz Initiative (ZT-0027) and the Christiane Nüsslein-Volhard Foundation fellowship. D.P. and M.L. were supported by the Helmholtz Graduate School for Cancer Research fellowship. H.-R.R. and T.H. were supported by Sonderforschungsbereich (SFB 873-B11) and H.-R.R. was supported by the European Research Council Advanced Grant 742883 and the Leibniz program of the Deutsche Forschungsgemeinschaft.

## Author contributions

L.F. and D.P. conceived the study, designed and performed experiments, interpreted results, wrote the original draft, and reviewed and edited the paper. M.L. developed the mathematical model, provided support with processing raw sequencing data, wrote, reviewed, and edited the paper. H.-

R.R. and T.H. supervised the study, wrote, reviewed and edited the paper. The manuscript was approved by all authors.

## Competing interests

The authors declare no competing interests.

## Data and materials availability

The data sets generated and/or analyzed during the current study are available from the corresponding authors upon reasonable request.

---

**Fig. S1.**
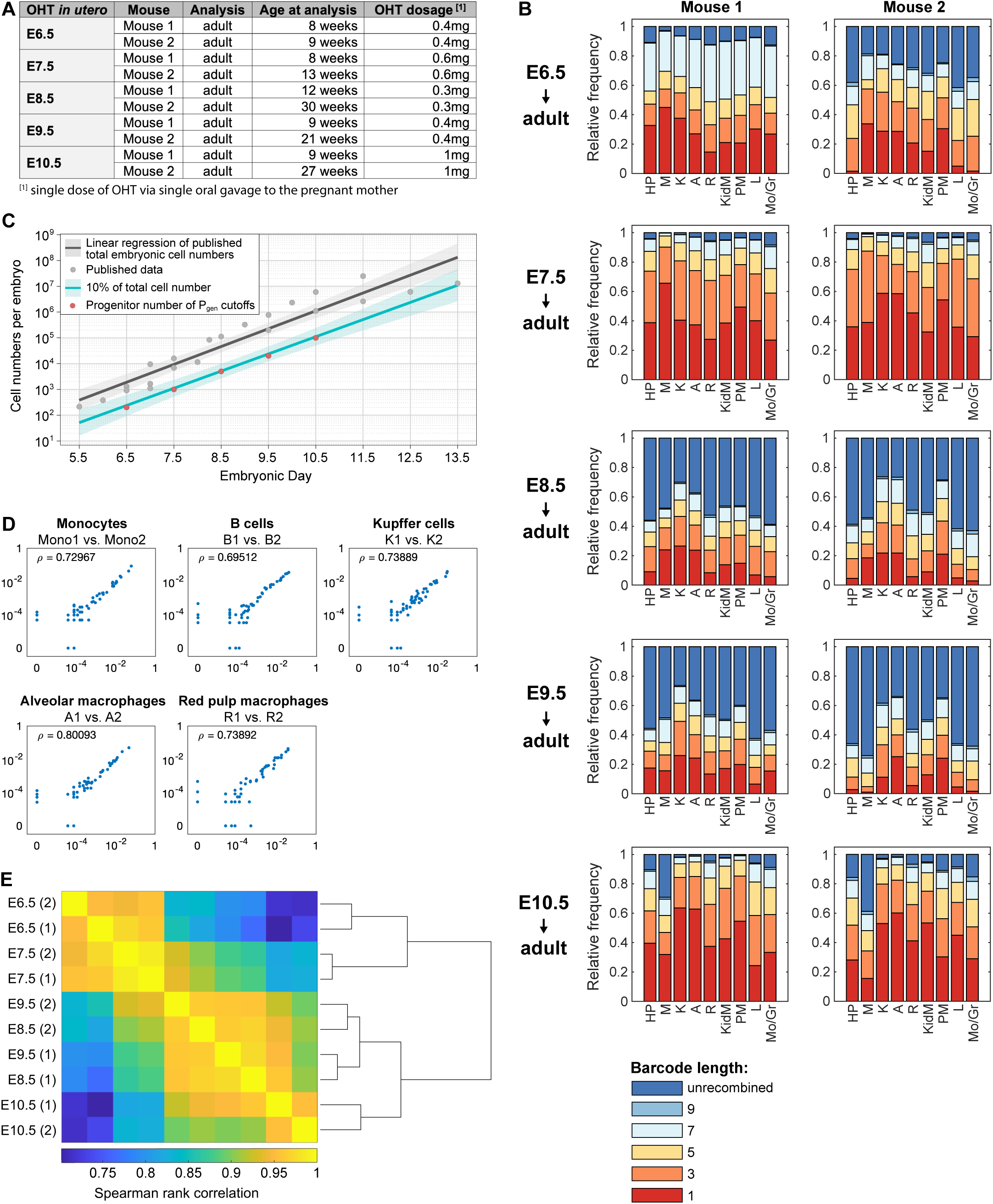
*Polylox* recombination patterns of individual *Rosa^Polylox/CreERT2^* mice and *P_gen_* cutoff estimation. (**A**) Table of analyzed mice with information on the age at analysis (in weeks) and OHT dosage that was applied as single oral gavage to the pregnant mother. (**B**) Recombination pattern of individual OHT-treated mice induced in utero at different timepoints (E6.5 – E10.5) as relative frequency for different sampled populations. Displayed as unrecombined (dark blue) or barcode length 9 (mid blue), 7 (light blue), 5 (yellow), 3 (orange), and 1 (red). n = 2 for each OHT timepoint *in utero*. Abbreviations for sampled populations: MG, microglia; KC, Kupffer cells; AM, Alveolar macrophages; KM, Kidney macrophages; PM, Peritoneal macrophages; RPM, Red pulp macrophages; L, Lymphocytes; Mo/Gr, Monocytes/Granulocytes; HP, hematopoietic stem and progenitor cells. (**C**) Estimated total embryonic cell number during development and progenitor number of selected *P_gen_* cutoffs. Linear regression of published data (***25–28***) of total embryonic cell counts, grey dots and dark grey line. As a conservative estimate of the potential number of macrophage progenitors, 10% of all total embryonic cells were chosen as upper limit, red line, and were used to determine suitable *P_gen_* cutoffs for each embryonic barcoding timepoint. (**D**) Correlation of sample repeats. Representative dot plots of different sorted populations showing overlap of sample repeats. Each dot represents an individual barcode. X- and y-axis show barcode frequencies. (**E**) Hierarchical meta clustering of clustered data sets of all analyzed mice using Spearman rank correlations of all sorted populations of each mouse. Mouse numbers (1, 2) indicate individual mice analyzed per OHT-induction timepoint.

**Fig S2:**
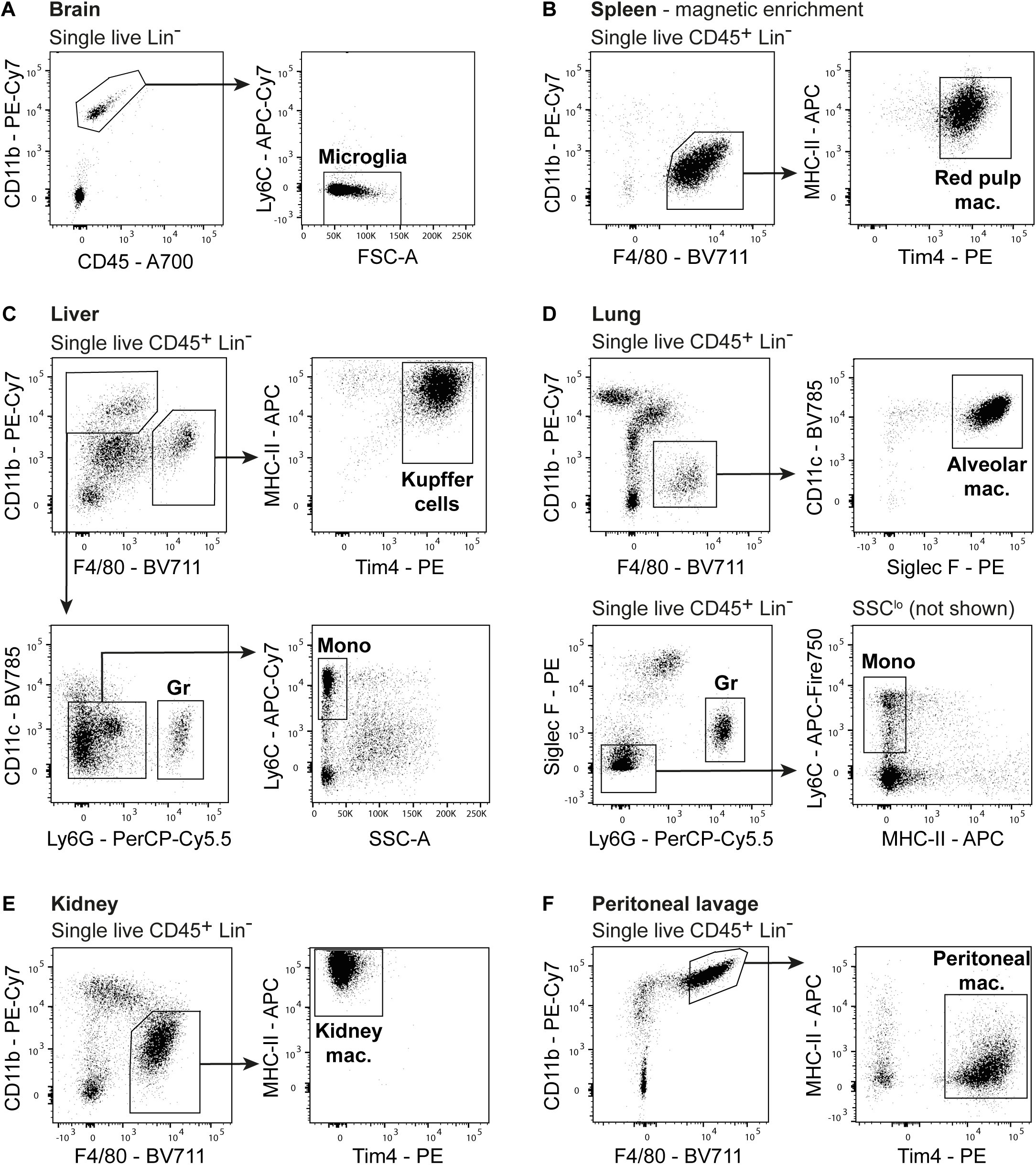
Gating schemes of tissue macrophages and myeloid populations for fluorescence-activated cell sorting. (**A**) Gating strategy for microglia (MG) in the brain, (**B**) red pulp macrophages (RPM) in the spleen, (**C**) Kupffer cells (KC), granulocytes (Gr), and monocytes (Mo) in the liver, (**D**) alveolar macrophages (AM), granulocytes (Gr), and monocytes (Mo) in the lung, (**E**) kidney macrophages (KM), (**F**) peritoneal macrophages (PM) from peritoneal exudate cells. Representative examples from an adult OHT-treated *Rosa^Polylox/CreERT2^* mouse at the age of eight weeks.

**Fig. S3:**
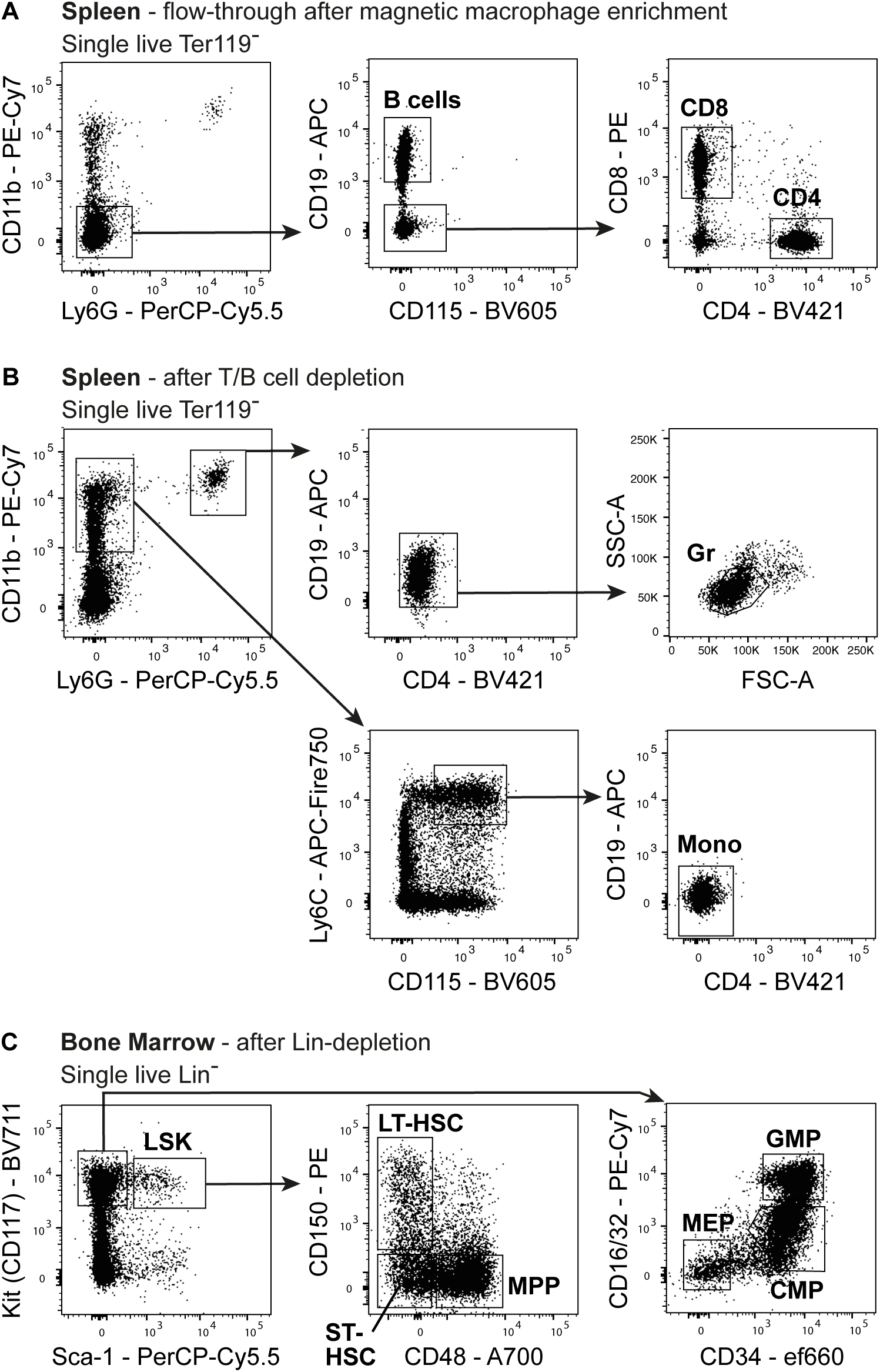
Gating schemes of lymphoid and myeloid populations from the spleen and hematopoietic stem and progenitor cells from the bone marrow for fluorescence-activated cell sorting. (**A**) Gating strategy for B cells, CD8, and CD4 T cells in the spleen sorted from splenic cells after magnetic-based isolation of red pulp macrophages. (**B**) Splenic monocytes (Mo) and granulocytes (Gr) after T and B cell depletion. (**C**) LSK (Lin^-^Sca-1^+^Kit^+^), long-term (LT-HSC) and short-term HSC (ST-HSC), multipotent progenitors (MPP), megakaryocyte-erythroid progenitor (MEP), common myeloid progenitor (CMP), and granulocyte-macrophage progenitor (GMP) sorted from Lin-depleted bone marrow cells. Representative examples from an adult OHT-treated *Rosa^Polylox/CreERT2^* mouse at the age of eight weeks.

**Fig. S4.**
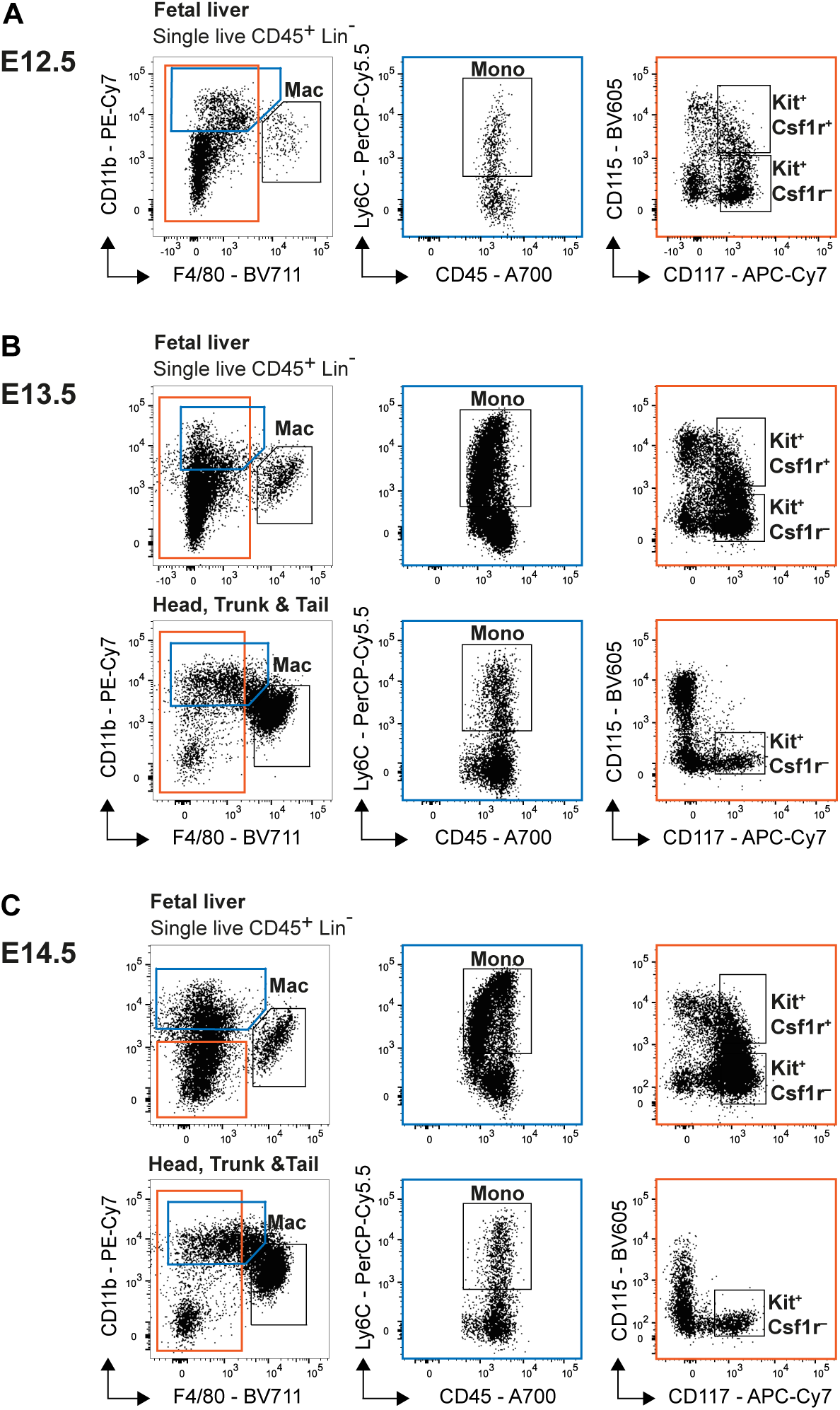
Gating schemes of macrophages, monocytes and progenitor populations isolated from fetuses for fluorescence-activated cell sorting. CD11b^+^F4/80^+^ macrophages (Mac), CD11b^+^Ly6C^+^ monocytes (Mono), and F4/80^-^Kit^+^ progenitors were sorted from the fetal liver, head, trunk, and tail region. Kit^+^Csf1r(CD115)^+^ progenitors were only sorted from the fetal liver from fetuses analyzed (**A**) at E12.5, (**B**) E13.5, and (**C**) E14.5.

**Fig. S5.**
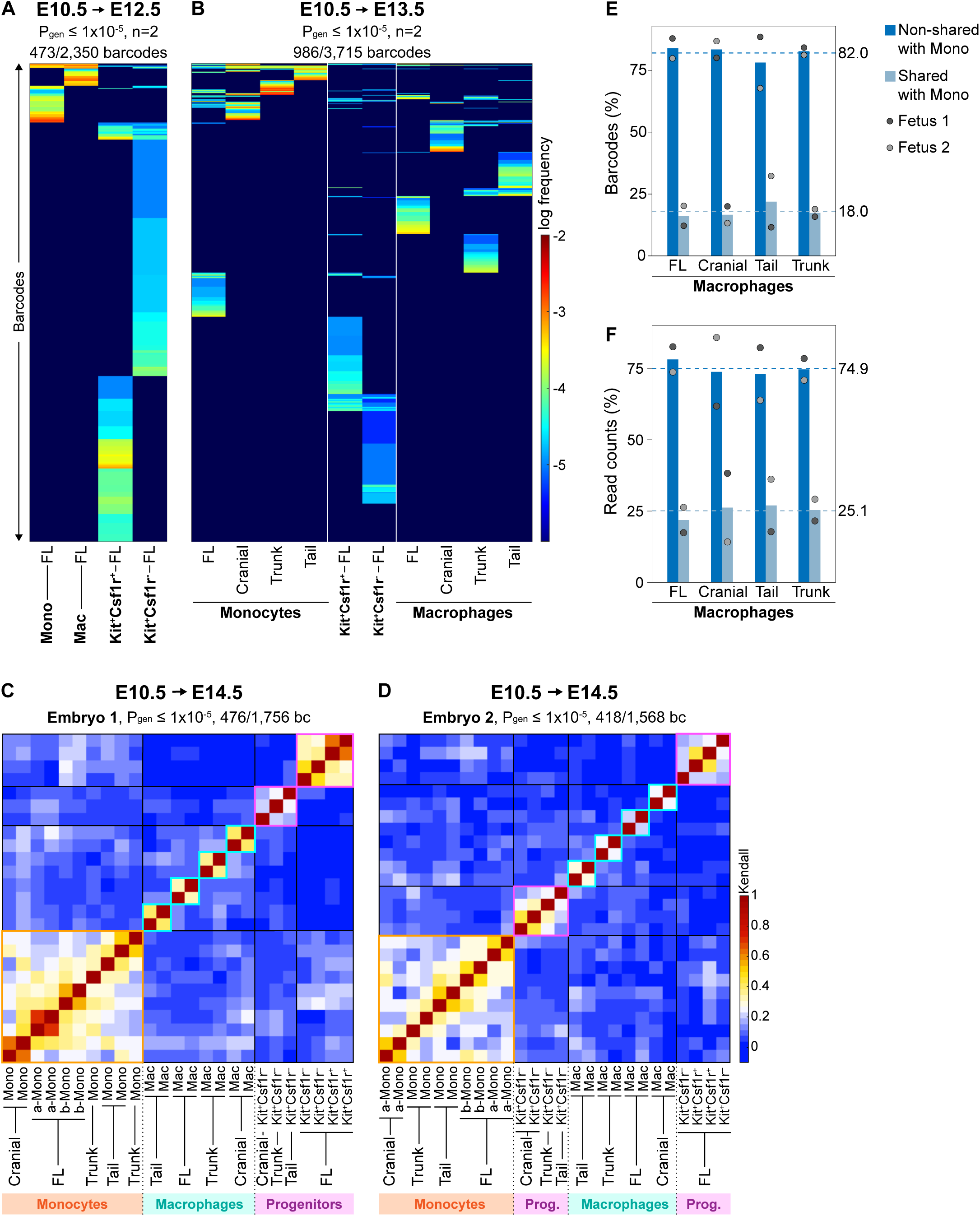
Extended data of fetuses analyzed at E12.5, E13.5 and E14.5. (**A-B**) Heatmaps of *Rosa^Polylox/CreERT2^*fetuses barcoded by a single oral gavage of OHT at E10.5 and analyzed at (**A**) E12.5 and (**B**) E13.5 (n=2 per timepoint) showing all filtered barcodes (*P_gen_* cutoff of ≤ 10^-5^) of populations from different fetal regions (macrophages, monocytes, progenitors). Numbers of total barcodes/*P_gen_* filtered barcodes are provided above each plot (**C-D**) Sample correlation heatmap of individual fetuses analyzed E14.5 with hierarchical clustering of all analyzed samples and annotated in major branches of hierarchical Kendall rank clustering as monocytes (orange), macrophages (turquoise), progenitors (pink). a-mono = Ly6C^+^CD115^-^ monocytes, b-mono = Ly6C^+^CD115^+^ monocytes, Mac = macrophages, bc = barcodes, FL = fetal liver. (**E**) Comparison of shared and non-shared barcode numbers (**F**) and read-based comparisons of non-shared barcodes (in %) of fetal monocytes and macrophages from different fetal fragments.

**Fig. S6.**
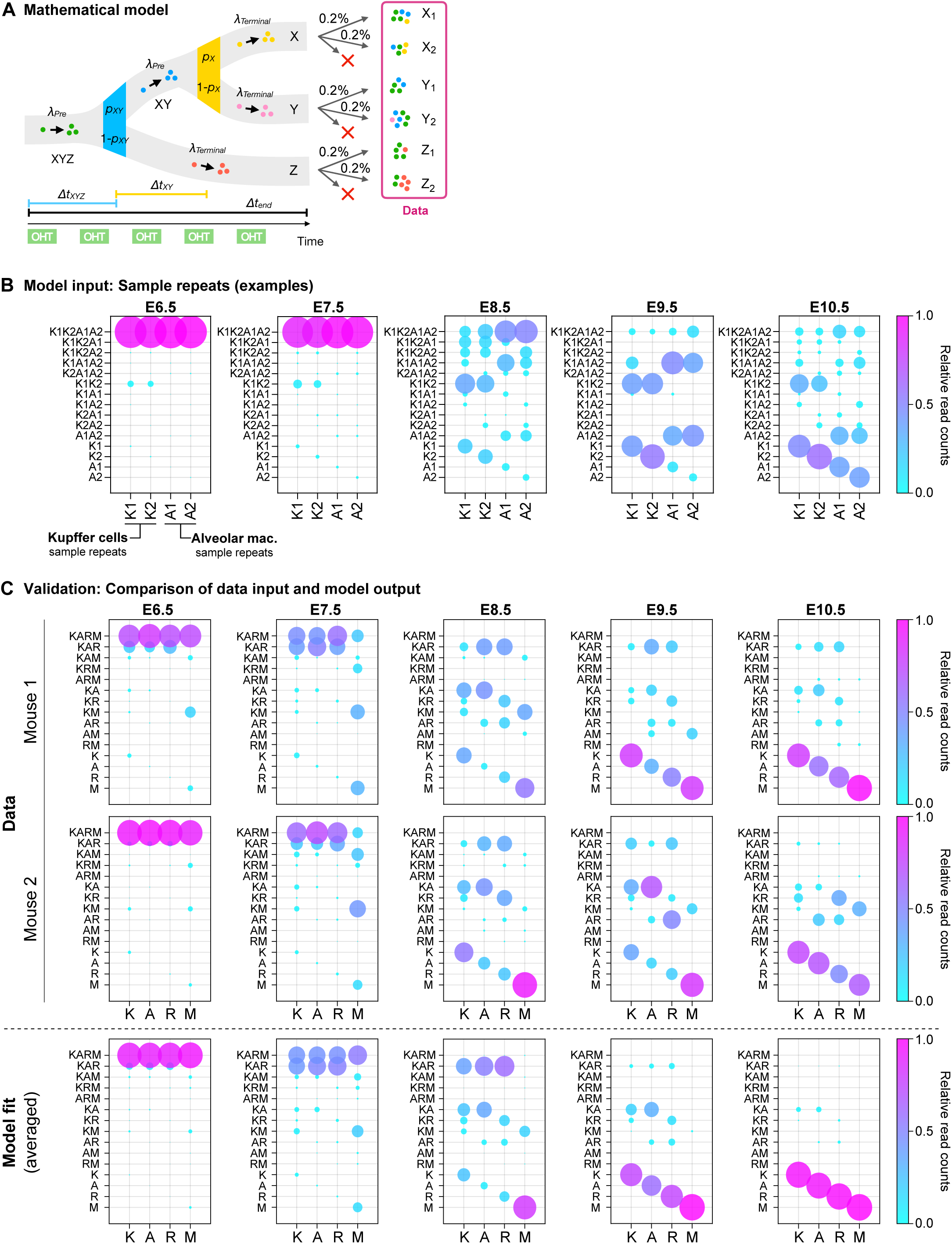
Extended data for modeling embryonic macrophage development using *Polylox* data. (**A**) Framework of the implemented generative mathematical model where sample repeats (biological replicates of one population from an individual mouse), considering their sampling efficiency, are compared for read-based barcode sharing to infer the probabilities of lineage bias at restriction events (*p*_AB_, 1-*p*_AB_ and *p*_A_, 1-*p*_A_), time until lineage restrictions (Δ*t_ABC_*, Δ*t_AB_*), time until final population is generated (Δ*t_end_*) as well as division rates of shared progenitor stages (*λ_Pre_*) and terminal unilineage progenitors (*λ_Terminal_*). (**B**) Exemplary sample repeats of Kupffer cells and Alveolar macrophages used as input data for the mathematical model. (**C**) Comparison of bubble plots of experimental data from individual mice (upper and middle row) and the average result from the model (lower row); sample repeats merged. Data used for modeling contains up to eight sample repeats and 255 fate combinations per mouse.

**Fig. S7.**
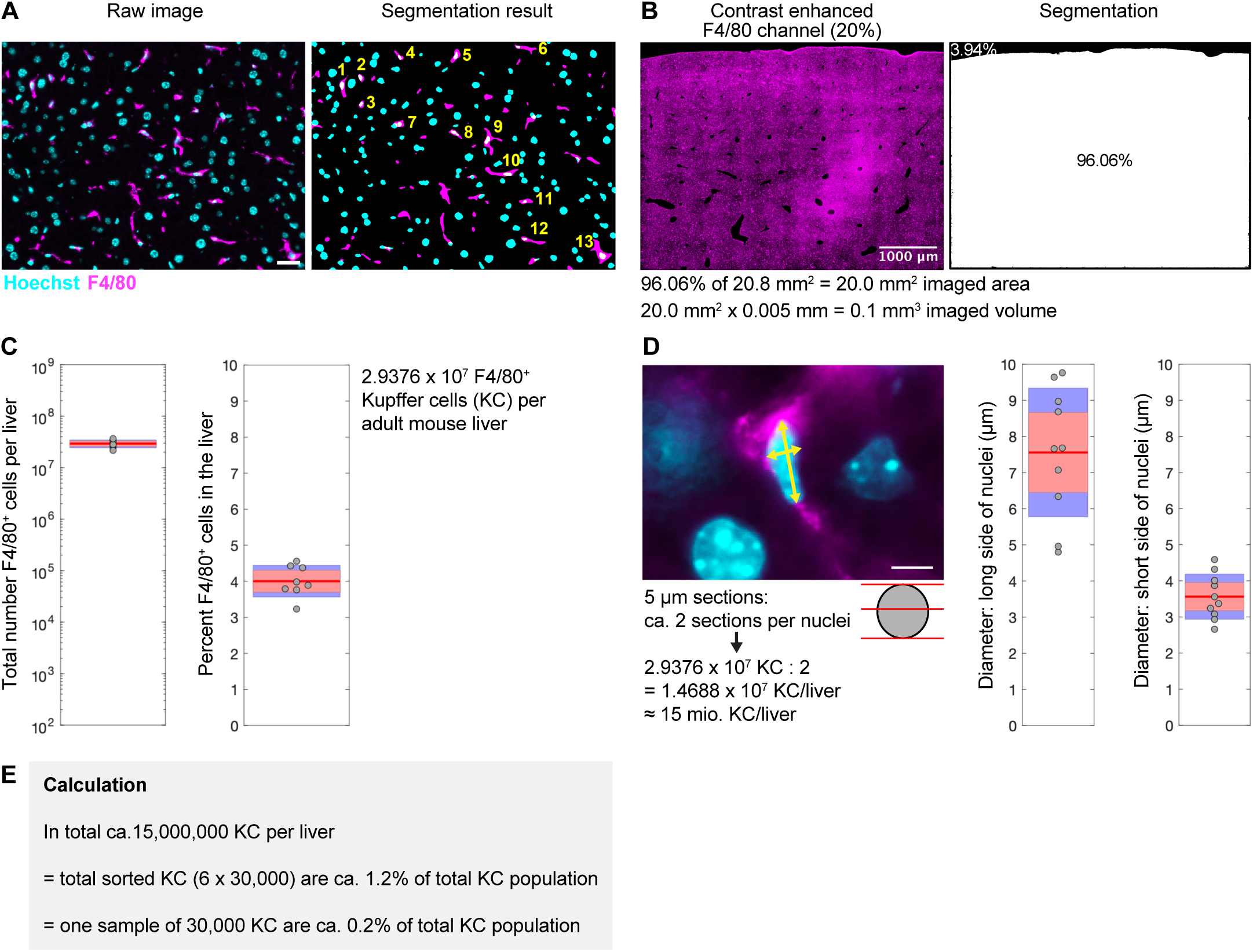
Sample loss estimation of total Kupffer cell population by isolation and sorting. (**A**) Raw and segmented images of immunofluorescent microscopy of 5µm liver sections stained with anti-F4/80 (magenta) and Hoechst (turquoise) nuclei staining. Scale bar shows 25 µm. (**B**) Segmentation of liver tissue and subtraction of non-liver tissue areas. (**C**) Total number of F4/80^+^ cells per liver (left) and percent of F4/80^+^ cells per liver (right). (**D**) Measurement of Kupffer cell nuclei diameters to correct for double detection of single macrophage nuclei that were potentially sectioned twice. Scale bar shows 5 µm. (**E**) Calculation of the Kupffer cell sample size (isolated for barcoding experiments) as a fraction of all Kupffer cells per liver.

**Fig. S8.**
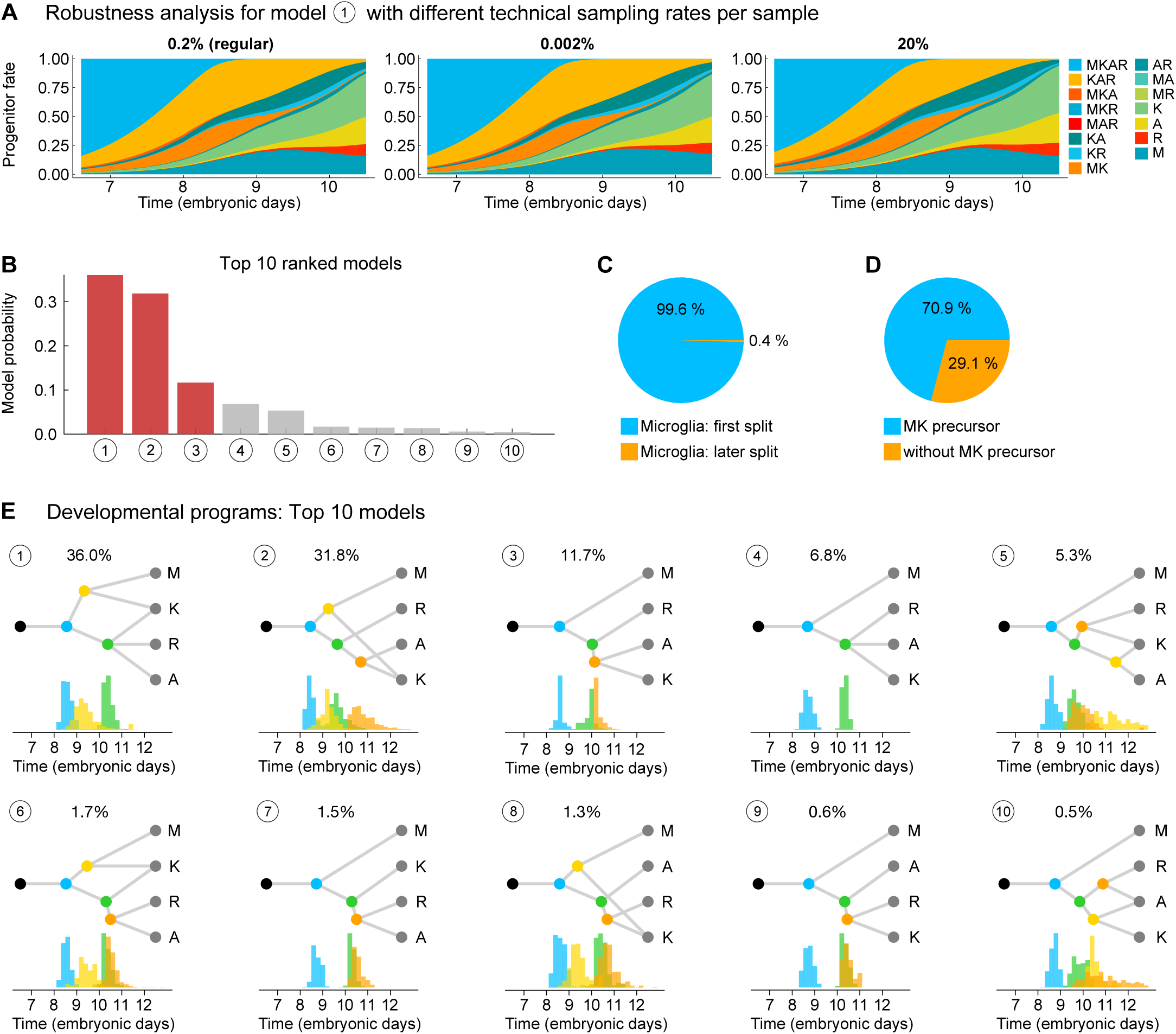
Extended data of inferred fates of embryonic macrophage progenitors from modeling *Polylox* data. (**A**) Robustness of model was tested with progenitor fate map by altering sampling efficiency. (**B**) Inferred probabilities of the top 10 ranked models of all 86 tested models, top 3 models in red. (**C**) Quantification of all models (weighted by model probability) with microglia splitting first (blue) and microglia splitting later (yellow). (**D**) Quantification of all models (weighted by model probability) with (blue) and without a MK progenitor (yellow). (**E**) Top 10 ranked models and their developmental programs with indicated model probabilities (percentage on top) and inferred branching times.

**Fig. S9.**
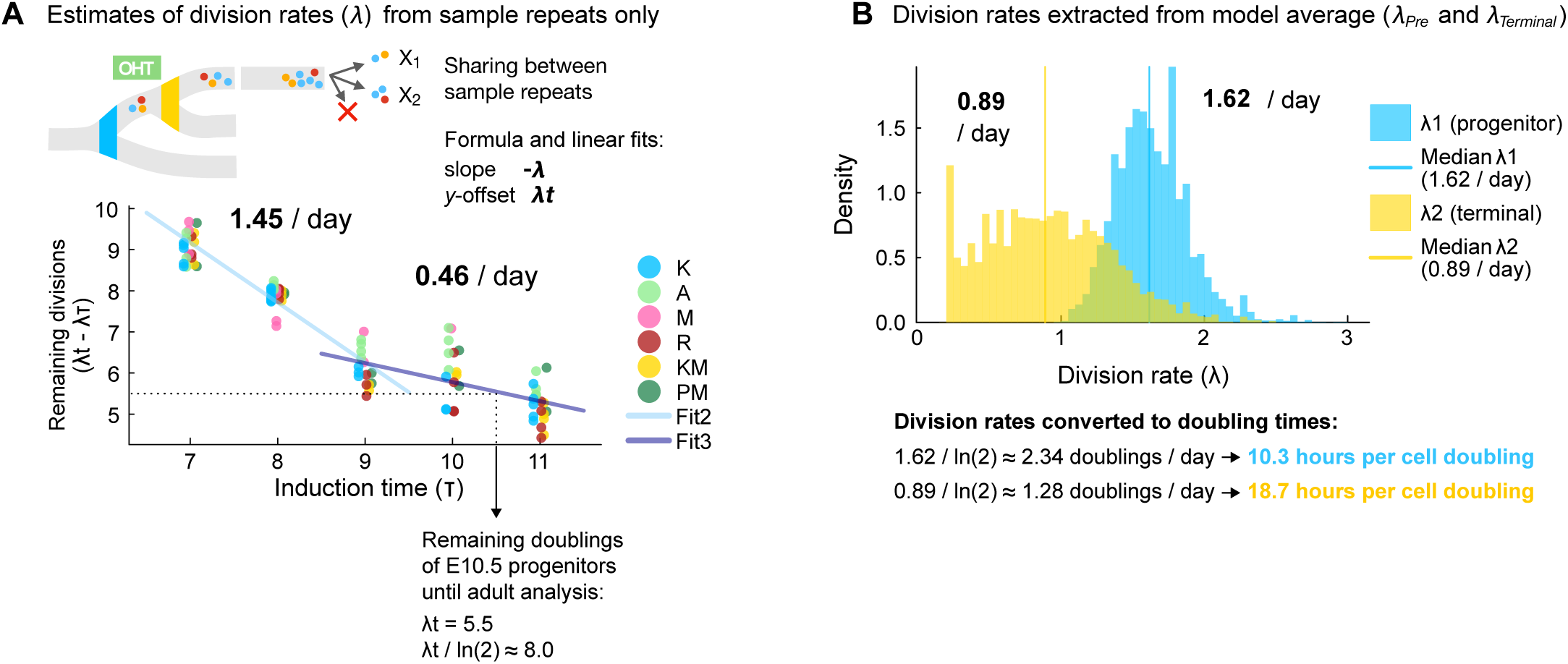
Inferred progenitor proliferation using two different approaches. (**A**) Estimates of division rates (*λ*) from sample repeats for different tissue macrophage populations with calculation to infer remaining doublings of progenitors at E10.5. (**B**) Division rates inferred from combined model average before (*λ_Pre_*) and after terminal lineage segregation (*λ_Terminal_*) and calculated doubling times of the populations.

**Fig. S10.**
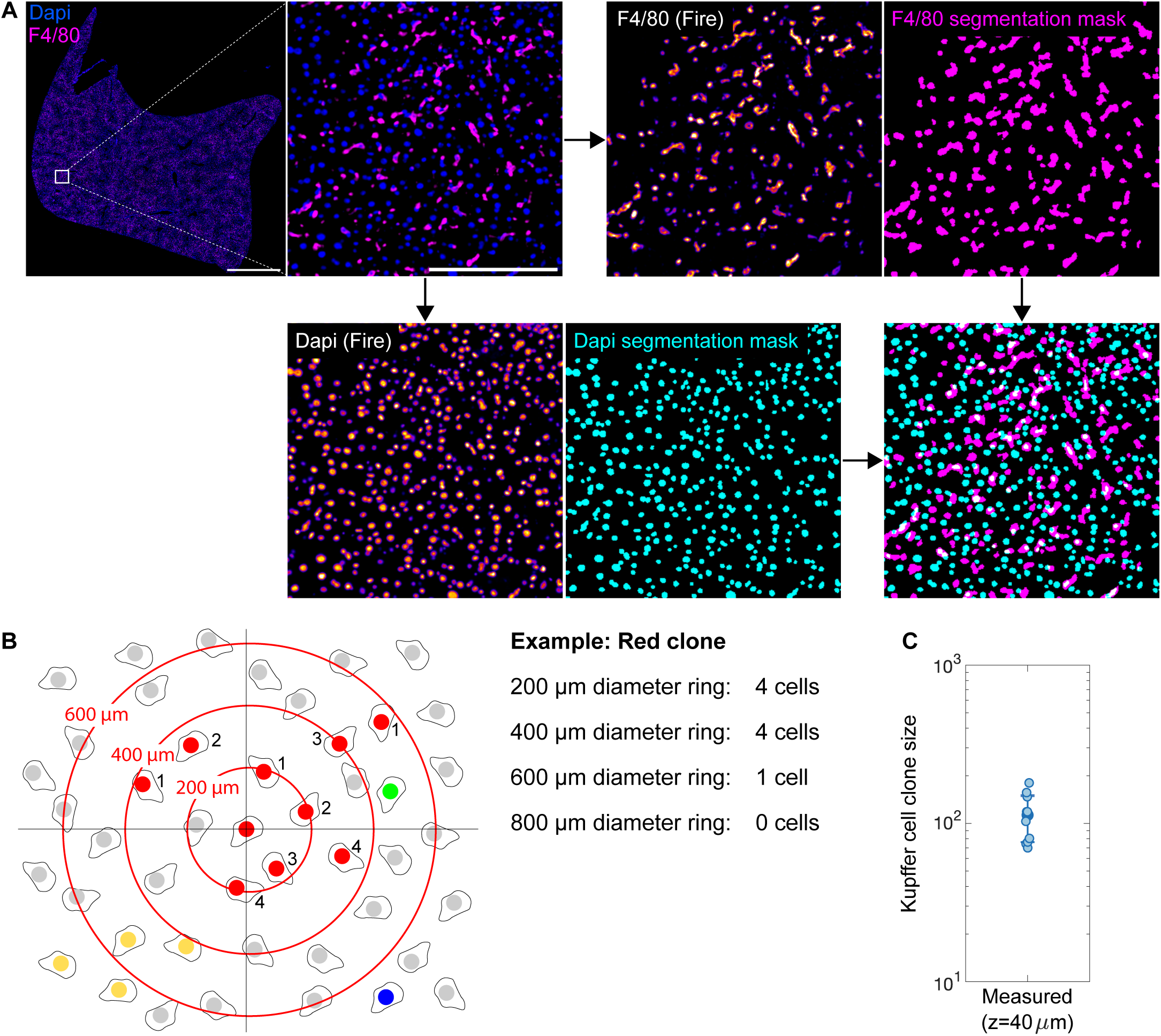
Spatial fate-mapping of Kupffer-cell progenitors with *Polytope* epitope barcoding. (**A**) Whole cryosections (5 µm) of adult *Rosa^Polytope/CreERT2^* mouse livers were analyzed. Depicted are segmentation results for nuclei and Kupffer cells (F4/80) Scale: 500 µm. (**B**) Approach for detecting spatial patterns of Kupffer cell clones. Concentric rings are drawn around the central cell and the number of cells with same color code is counted. (**C**) Measured Kupffer cell clone sizes from in total 40 µm in z.

**Fig. S11.**
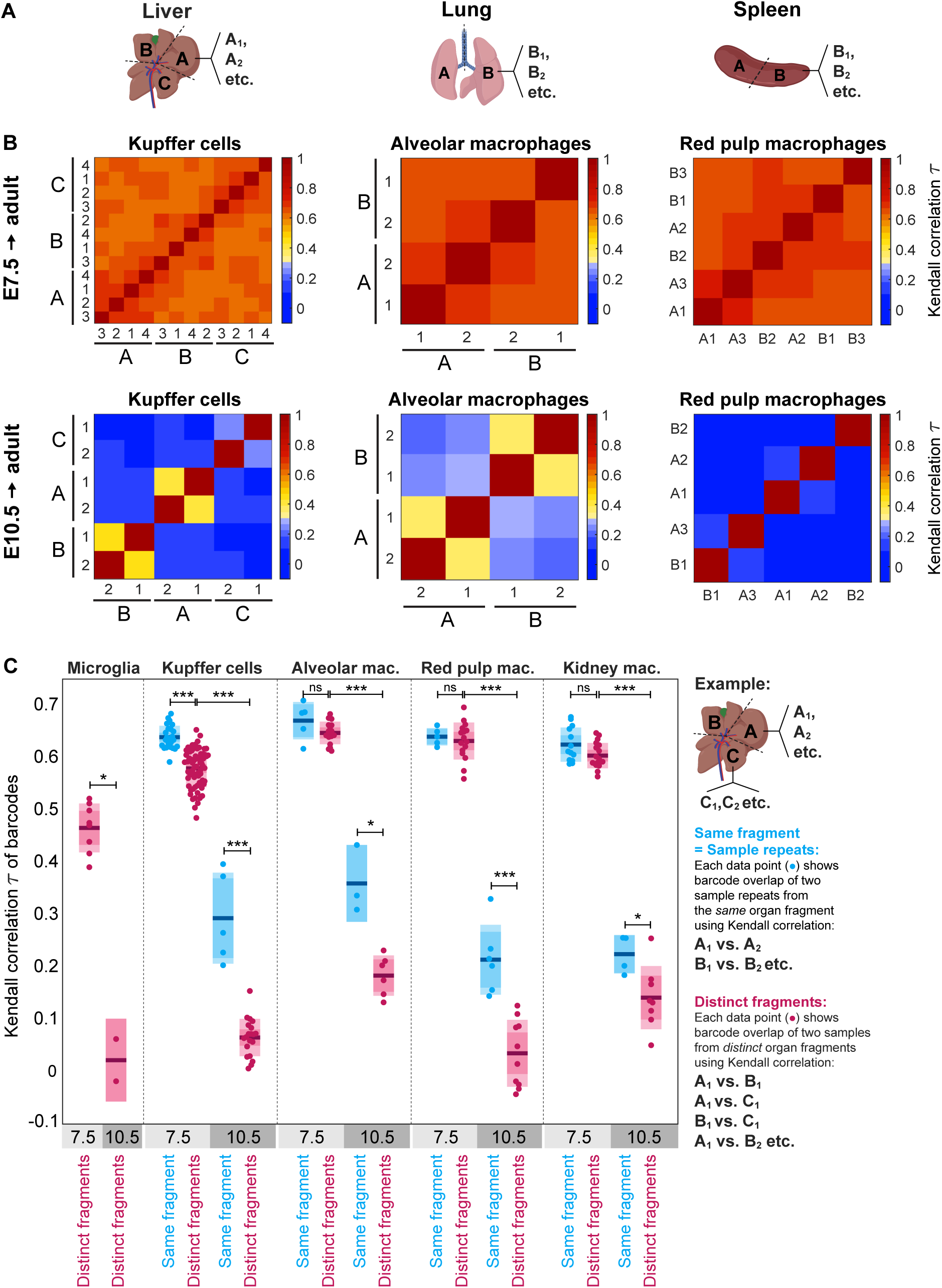
Comparison of Kendall rank correlation of barcodes from sample repeats of distinct anatomical sites. (**A**) Analysis of sample repeats from the same and distinct organ fragments of liver, lung, and spleen. (**B**) Sample repeats of distinct liver lobes (A, B, and C), left and right lobe of the lung (A and B), and spleen halves (A, B) at E7.5 (upper row) and at E10.5 (lower row). (**C**) Box plot of sample correlation using Kendall‘s tau correlation values when comparing samples from distinct anatomical fragments (red) or sample repeats from the same anatomical fragment (blue). Box plots show the mean, 1σ, and the 95% confidence interval. Each individual dot represents one comparison between the same fragment (blue) or distinct fragments (red). Stars indicate statistically significant values (Wilcoxon-rank testing) with p-values ≤ 0.05 (*) and ≤ 0.001 (***), > 0.05 as not significant (ns), all p-values are listed in Table S4.

**Table S1:** Numbers of barcode contributions (relative read counts) to sampled populations from adult analyzed mice (Fig. 1C, E, G, I, K). H, hematopoietic progenitors; M, microglia; K, Kupffer cells; A, alveolar macrophages; R, red pulp macrophages. Turquoise: common mult-ilineage progenitor HMKAR; pink: progenitor with hematopoietic and partial macrophage potential (HKAR, HKR, HR, HA, HK). Green: macrophage progenitor without microglia fates (KAR, AR, KR, KA); yellow: bilineage microglia-Kupffer cell progenitor (MK); blue: unilineage progenitors (H, M, K, A, R).

|  | E6.5 → adult |  |  |  |  | E7.5 → adult |  |  |  |  | E8.5 → adult |  |  |  |  |
| --- | --- | --- | --- | --- | --- | --- | --- | --- | --- | --- | --- | --- | --- | --- | --- |
| Fates | H | M | K | A | R | H | M | K | A | R | H | M | K | A | R |
| HMKAR | 90% | 78% | 79% | 85% | 81% | 68% | 14% | 57% | 60% | 62% | 0% | 0% | 0% | 0% | 0% |
| MKAR | 0% | 1% | 1% | 0% | 0% | 0% | 10% | 14% | 13% | 9% | 0% | 0% | 0% | 0% | 1% |
| HKAR | 8% | 0% | 13% | 13% | 17% | 27% | 0% | 14% | 20% | 24% | 9% | 0% | 0% | 0% | 1% |
| HMAR | 0% | 0% | 0% | 0% | 0% | 0% | 0% | 0% | 0% | 0% | 0% | 0% | 0% | 0% | 0% |
| HMKR | 0% | 2% | 0% | 0% | 0% | 0% | 2% | 0% | 0% | 0% | 3% | 0% | 0% | 0% | 1% |
| HMKA | 0% | 0% | 0% | 0% | 0% | 0% | 2% | 1% | 0% | 0% | 0% | 0% | 0% | 0% | 0% |
| KAR | 0% | 0% | 1% | 1% | 1% | 0% | 0% | 4% | 4% | 4% | 0% | 0% | 16% | 50% | 40% |
| MAR | 0% | 0% | 0% | 0% | 0% | 0% | 0% | 0% | 0% | 0% | 0% | 0% | 0% | 0% | 0% |
| MKR | 0% | 1% | 0% | 0% | 0% | 0% | 11% | 1% | 0% | 0% | 0% | 0% | 0% | 0% | 0% |
| MKA | 0% | 4% | 2% | 0% | 0% | 0% | 8% | 2% | 0% | 0% | 0% | 4% | 1% | 0% | 0% |
| HAR | 0% | 0% | 0% | 0% | 0% | 0% | 0% | 0% | 0% | 0% | 0% | 0% | 0% | 0% | 0% |
| HKR | 0% | 0% | 0% | 0% | 0% | 3% | 0% | 0% | 0% | 1% | 67% | 0% | 2% | 0% | 6% |
| HKA | 0% | 0% | 0% | 0% | 0% | 1% | 0% | 1% | 0% | 0% | 0% | 0% | 0% | 0% | 0% |
| HMR | 0% | 0% | 0% | 0% | 0% | 0% | 0% | 0% | 0% | 0% | 0% | 0% | 0% | 0% | 0% |
| HMA | 0% | 0% | 0% | 0% | 0% | 0% | 0% | 0% | 0% | 0% | 0% | 0% | 0% | 0% | 0% |
| HMK | 0% | 5% | 0% | 0% | 0% | 0% | 3% | 0% | 0% | 0% | 2% | 0% | 0% | 0% | 0% |
| AR | 0% | 0% | 0% | 0% | 0% | 0% | 0% | 0% | 0% | 0% | 0% | 0% | 0% | 7% | 13% |
| KR | 0% | 0% | 0% | 0% | 0% | 0% | 0% | 0% | 0% | 0% | 0% | 0% | 9% | 0% | 23% |
| KA | 0% | 0% | 1% | 0% | 0% | 0% | 0% | 0% | 0% | 0% | 0% | 0% | 28% | 32% | 0% |
| MR | 0% | 0% | 0% | 0% | 0% | 0% | 0% | 0% | 0% | 0% | 0% | 2% | 0% | 0% | 0% |
| MA | 0% | 0% | 0% | 0% | 0% | 0% | 0% | 0% | 0% | 0% | 0% | 1% | 0% | 0% | 0% |
| MK | 0% | 9% | 1% | 0% | 0% | 0% | 46% | 3% | 0% | 0% | 0% | 24% | 3% | 0% | 0% |
| HR | 0% | 0% | 0% | 0% | 0% | 0% | 0% | 0% | 0% | 0% | 0% | 0% | 0% | 0% | 0% |
| HA | 2% | 0% | 0% | 0% | 0% | 0% | 0% | 0% | 0% | 0% | 7% | 0% | 0% | 0% | 0% |
| HK | 0% | 0% | 1% | 0% | 0% | 0% | 0% | 1% | 0% | 0% | 3% | 0% | 0% | 0% | 0% |
| HM | 0% | 0% | 0% | 0% | 0% | 0% | 0% | 0% | 0% | 0% | 0% | 0% | 0% | 0% | 0% |
| R | 0% | 0% | 0% | 0% | 0% | 0% | 0% | 0% | 0% | 0% | 0% | 0% | 0% | 0% | 14% |
| A | 0% | 0% | 0% | 0% | 0% | 0% | 0% | 0% | 0% | 0% | 0% | 0% | 0% | 10% | 0% |
| K | 0% | 0% | 1% | 0% | 0% | 0% | 0% | 2% | 0% | 0% | 0% | 0% | 39% | 0% | 0% |
| M | 0% | 1% | 0% | 0% | 0% | 0% | 4% | 0% | 0% | 0% | 0% | 69% | 0% | 0% | 0% |
| H | 0% | 0% | 0% | 0% | 0% | 0% | 0% | 0% | 0% | 0% | 9% | 0% | 0% | 0% | 0% |

(Continuation of Table S1)
|  | E9.5 → adult |  |  |  |  | E10.5 → adult |  |  |  |  |
| --- | --- | --- | --- | --- | --- | --- | --- | --- | --- | --- |
| Fates | H | M | K | A | R | H | M | K | A | R |
| HMKAR | 0% | 0% | 0% | 0% | 0% | 0% | 0% | 0% | 0% | 0% |
| MKAR | 0% | 0% | 0% | 0% | 0% | 0% | 1% | 0% | 0% | 0% |
| HKAR | 0% | 0% | 0% | 0% | 1% | 3% | 0% | 0% | 0% | 0% |
| HMAR | 0% | 0% | 0% | 0% | 0% | 0% | 0% | 0% | 0% | 0% |
| HMKR | 0% | 0% | 0% | 0% | 0% | 0% | 0% | 0% | 0% | 0% |
| HMKA | 0% | 0% | 0% | 0% | 0% | 0% | 0% | 0% | 0% | 0% |
| KAR | 0% | 0% | 27% | 58% | 55% | 0% | 0% | 10% | 27% | 15% |
| MAR | 0% | 0% | 0% | 0% | 0% | 0% | 0% | 0% | 0% | 0% |
| MKR | 0% | 0% | 0% | 0% | 0% | 0% | 0% | 0% | 0% | 0% |
| MKA | 0% | 0% | 0% | 0% | 0% | 0% | 2% | 0% | 0% | 0% |
| HAR | 0% | 0% | 0% | 0% | 1% | 4% | 0% | 0% | 1% | 4% |
| HKR | 59% | 0% | 0% | 0% | 2% | 0% | 0% | 0% | 0% | 0% |
| HKA | 0% | 0% | 0% | 0% | 0% | 0% | 0% | 0% | 0% | 0% |
| HMR | 0% | 0% | 0% | 0% | 0% | 3% | 2% | 0% | 0% | 0% |
| HMA | 0% | 0% | 0% | 0% | 0% | 0% | 0% | 0% | 0% | 0% |
| HMK | 0% | 0% | 0% | 0% | 0% | 0% | 0% | 0% | 0% | 0% |
| AR | 0% | 0% | 0% | 9% | 12% | 0% | 0% | 0% | 10% | 9% |
| KR | 0% | 0% | 4% | 0% | 14% | 0% | 0% | 9% | 0% | 19% |
| KA | 0% | 0% | 15% | 15% | 0% | 0% | 0% | 14% | 15% | 0% |
| MR | 0% | 0% | 0% | 0% | 0% | 0% | 0% | 0% | 0% | 0% |
| MA | 0% | 11% | 0% | 2% | 0% | 0% | 0% | 0% | 0% | 0% |
| MK | 0% | 34% | 1% | 0% | 0% | 0% | 24% | 1% | 0% | 0% |
| HR | 31% | 0% | 0% | 0% | 1% | 10% | 0% | 0% | 0% | 5% |
| HA | 3% | 0% | 0% | 0% | 0% | 9% | 0% | 0% | 1% | 0% |
| HK | 4% | 0% | 1% | 0% | 0% | 0% | 0% | 0% | 0% | 0% |
| HM | 0% | 0% | 0% | 0% | 0% | 3% | 0% | 0% | 0% | 0% |
| R | 0% | 0% | 0% | 0% | 14% | 0% | 0% | 0% | 0% | 46% |
| A | 0% | 0% | 0% | 14% | 0% | 0% | 0% | 0% | 47% | 0% |
| K | 0% | 0% | 51% | 0% | 0% | 0% | 0% | 65% | 0% | 0% |
| M | 0% | 55% | 0% | 0% | 0% | 0% | 71% | 0% | 0% | 0% |
| H | 2% | 0% | 0% | 0% | 0% | 68% | 0% | 0% | 0% | 0% |

**Table S2:** Numbers of barcode contributions (relative read counts) to sampled populations from E14.5 analyzed fetuses (Fig. 2D). M/Monos, monocytes; P/Kit^+^ Prog., CD45^+^Lin^-^Kit^+^ progenitors; L, fetal liver macrophages; C, cranial macrophages; T, tail/trunk macrophages. Green: progenitors with Kit^+^ progenitor and macrophage fates; yellow: progenitors with monocyte and macrophage fates; blue: unilineage fates.

| Fates | Monos<br>M | Kit<sup>+</sup> Prog.<br>P | Macrophage |  |  |
| --- | --- | --- | --- | --- | --- |
|  |  |  | Fetal liver<br>L | Cranial<br>C | Tail/trunk<br>T |
| **MPLCT** | 0,3% | 0,1% | 1,3% | 0,7% | 0,7% |
| **PLCT** | 0,0% | 0,0% | 0,0% | 0,0% | 0,0% |
| **MLCT** | 1,5% | 0,0% | 4,5% | 11,9% | 4,4% |
| **MPCT** | 0,9% | 0,1% | 0,0% | 2,4% | 1,3% |
| **MPLT** | 0,0% | 0,0% | 0,0% | 0,0% | 0,0% |
| **MPLC** | 0,0% | 0,0% | 0,0% | 0,0% | 0,0% |
| **LCT** | 0,0% | 0,0% | 1,0% | 0,8% | 0,5% |
| **PCT** | 0,0% | 0,1% | 0,0% | 0,4% | 0,1% |
| **PLT** | 0,0% | 1,4% | 0,1% | 0,0% | 0,1% |
| **PLC** | 0,0% | 1,6% | 0,9% | 0,5% | 0,0% |
| **MCT** | 2,0% | 0,0% | 0,0% | 5,6% | 2,9% |
| **MLT** | 5,0% | 0,0% | 4,3% | 0,0% | 7,1% |
| **MLC** | 0,0% | 0,0% | 0,3% | 0,6% | 0,0% |
| **MPT** | 3,2% | 2,2% | 0,0% | 0,0% | 0,8% |
| **MPC** | 1,3% | 0,9% | 0,0% | 1,7% | 0,0% |
| **MPL** | 0,7% | 2,1% | 4,9% | 0,0% | 0,0% |
| **CT** | 0,0% | 0,0% | 0,0% | 3,4% | 3,2% |
| **LT** | 0,0% | 0,0% | 10,6% | 0,0% | 7,5% |
| **LC** | 0,0% | 0,0% | 2,1% | 3,4% | 0,0% |
| **PT** | 0,0% | 0,8% | 0,0% | 0,0% | 1,3% |
| **PC** | 0,0% | 1,0% | 0,0% | 1,9% | 0,0% |
| **PL** | 0,0% | 4,0% | 4,3% | 0,0% | 0,0% |
| **MT** | 11,0% | 0,0% | 0,0% | 0,0% | 8,0% |
| **MC** | 5,0% | 0,0% | 0,0% | 4,3% | 0,0% |
| **ML** | 7,7% | 0,0% | 5,7% | 0,0% | 0,0% |
| **MP** | 22,5% | 21,9% | 0,0% | 0,0% | 0,0% |
| **T** | 0,0% | 0,0% | 0,0% | 0,0% | 62,3% |
| **C** | 0,0% | 0,0% | 0,0% | 62,5% | 0,0% |
| **L** | 0,0% | 0,0% | 60,1% | 0,0% | 0,0% |
| **P** | 0,0% | 63,7% | 0,0% | 0,0% | 0,0% |
| **M** | 38,8% | 0,0% | 0,0% | 0,0% | 0,0% |

**Table S3:** Progenitor estimates (Fig. 3I) and mean clone sizes of adult Kupffer cells (Fig. 3J) inferred from mathematical modeling.

| Timepoint | Kupffer cell progenitor estimates | Mean clone sizes of adult Kupffer cells |
| --- | --- | --- |
| E6.5 | 257 | 58,435 |
| E6.75 | 402 | 37,289 |
| E7.0 | 624 | 24,030 |
| E7.25 | 960 | 15,632 |
| E7.5 | 1,455 | 10,308 |
| E7.75 | 2,176 | 6,893 |
| E8.0 | 3,197 | 4,692 |
| E8.25 | 4,583 | 3,273 |
| E8.5 | 6,436 | 2,331 |
| E8.75 | 9,096 | 1,649 |
| E9.0 | 12,775 | 1,174 |
| E9.25 | 17,590 | 853 |
| E9.5 | 23,874 | 628 |
| E9.75 | 32,190 | 466 |
| E10.0 | 42,883 | 350 |
| E10.25 | 55,458 | 270 |
| E10.5 | 69,467 | 216 |
| E10.75 | 86,393 | 174 |
| E11.0 | 106,149 | 141 |
| E11.25 | 129,101 | 116 |
| E11.5 | 155,298 | 97 |
| E11.75 | 184,865 | 81 |
| E12.0 | 218,784 | 69 |
| E12.25 | 256,404 | 59 |

**Table S4.** Statistical test results (Fig. S11C). Statistical comparisons were performed using Wilcoxon-rank testing. P-values ≤ 0.05 were considered significant. Distinct fragments, i.e., comparison of samples from different anatomical sites; same fragment, i.e., comparison of sample repeats from the same anatomical sites; ns = not significant.)

| Population | Comparison |  | P-value |
| --- | --- | --- | --- |
| **Microglia** | E7.5 Distinct fragments | E10.5 Distinct fragments | $4.44 \times 10^{-2}$ |
| **Kupffer cells** | E7.5 Same fragment | E7.5 Distinct fragments | $4.44 \times 10^{-11}$ |
| | E7.5 Distinct fragments | E10.5 Distinct fragments | $1.33 \times 10^{-11}$ |
| | E10.5 Same fragment | E10.5 Distinct fragment | $7.71 \times 10^{-4}$ |
| **Alveolar macrophages** | E7.5 Same fragment | E7.5 Distinct fragments | $1.07 \times 10^{-1}$ (ns) |
| | E7.5 Distinct fragments | E10.5 Distinct fragments | $4.62 \times 10^{-4}$ |
| | E10.5 Same fragment | E10.5 Distinct fragment | $2.38 \times 10^{-2}$ |
| **Red pulp macrophages** | E7.5 Same fragment | E7.5 Distinct fragments | $7.10 \times 10^{-1}$ (ns) |
| | E7.5 Distinct fragments | E10.5 Distinct fragments | $2.79 \times 10^{-5}$ |
| | E10.5 Same fragment | E10.5 Distinct fragment | $2.50 \times 10^{-4}$ |
| **Kidney macrophages** | E7.5 Same fragment | E7.5 Distinct fragments | $8.16 \times 10^{-2}$ (ns) |
| | E7.5 Distinct fragments | E10.5 Distinct fragments | $7.12 \times 10^{-5}$ |
| | E10.5 Same fragment | E10.5 Distinct fragment | $1.62 \times 10^{-2}$ |

## Notes

### Competing Interest Statement

The authors have declared no competing interest.

